# Mis-localization of PBPs in *Staphylococcus aureus gdpP* mutant contributes to β-lactam resistance and surface protein cross-wall trafficking

**DOI:** 10.1101/2025.04.15.649054

**Authors:** Yaosheng Jia, Ran Zhang, Salvatore J. Scaffidi, Wenqi Yu

## Abstract

Mutations in *gdpP* are frequently associated with β-lactam resistance in *Staphylococcus aureus*. GdpP is a phosphodiesterase that degrades the second messenger c-di-AMP, which plays a key role in osmoregulation and antibiotic resistance in many Gram-positive bacteria. However, the mechanisms of β-lactam resistance remain unknown. We show here that deletion of *gdpP* disrupted penicillin-binding proteins (PBPs)-mediated deposition of YSIRK+ surface protein SpA to septal peptidoglycan (cross-wall). In contrast to their septal localization in WT staphylococcal cells, all four PBPs (PBP1-4) were drastically mis-localized as distinct single foci in Δ*gdpP.* The mis-localization of PBPs is attributed to c-di-AMP accumulation as overexpression of c-di-AMP synthetase *dacA* phenocopied Δ*gdpP.* In addition, Δ*gdpP* exhibited severe cell cycle retardation deficient in cell division initiation. The aberrant foci formation of PBPs altered the spatial distribution of cell wall synthesis but did not affect the overall cell wall cross-linking. The foci formation of PBPs correlated with β-lactam resistance: the foci gradually decreased with increased concentrations of penicillin. We concluded that the aberrant spatial distribution of PBPs and the cell division defects of Δ*gdpP* contribute to β-lactam resistance and YSIRK+ protein cross-wall trafficking.

## Introduction

*Staphylococcus aureus* is a Gram-positive bacterium that frequently colonizes human nares and skin (1). It is also one of the most important pathogenic bacteria, responsible for causing various infections, such as skin and soft tissue infections, pneumonia, endocarditis, osteomyelitis, and life-threatening sepsis (2–4). β-lactam antibiotics, including cephalosporin and oxacillin are frequently used for *S. aureus* infections in clinical settings (5, 6). However, the rise of antibiotic-resistant *S. aureus*, including community and hospital MRSA (methicillin-resistant *Staphylococcus aureus*), poses a significant challenge to these primary treatment methods (7, 8). β-lactam antibiotics inhibit the transpeptidase activity of penicillin-binding proteins (PBPs), which is essential for cell wall peptidoglycan (PG) cross-linking. Remarkably, *S. aureus* has evolved various resistance mechanisms against generations of β-lactam antibiotics over the years. β-lactamase encoded by BlaZ contributes to narrow-spectrum resistance by degrading β-lactam antibiotics (9). The presence of the *mecA* gene, which encodes penicillin-binding protein 2a (PBP2a), renders resistance to β-lactamase-resistant antibiotics in MRSA strains (10, 11). The recent discovery of *mecA*-negative MRSA strains reveals that mutations in the *gdpP* gene are also frequently associated with high-level β-lactam resistance in both clinical and laboratory isolates (12–24).

While the association of *gdpP* mutation with β-lactam resistance has been reported in several Gram-positive bacteria, including *S. aureus*, *B. subtilis*, *L. lactis*, *L. monocytogenes*, *S. pyogenes*, *S. agalactiae*, and *E. faecalis* (13, 25–34), the underlying mechanism is largely unknown. GdpP (GGDEF domain protein containing phosphodiesterase) in *S. aureus* is a close homolog to YybT in *B. subtilis* and Pde1 in *S. pneumoniae*, which is known to hydrolyze c-di-AMP, a second messenger widely found in bacteria (13, 35, 36). In. *S. aureus*, a gene named *dacA* encodes for a diadenylate cyclase that synthesizes c-di-AMP from two ATP molecules (37). C-di-AMP regulates multiple facets of bacterial physiology, such as cell volume, osmotic homeostasis, cell wall homeostasis, metabolism, DNA repair and virulence (13, 38–43). One of the conserved functions of c-di-AMP across bacterial species is osmoregulation by binding to proteins and riboswitches that regulate potassium and compatible solutes transportation (44, 45). In *S. aureus,* c-di-AMP has been shown to bind to KtrA (potassium transporter-gating component), CpaA (a predicted cation/proton antiporter), PstA (PII-like signal transduction protein A), KdpD (a sensor histidine kinase that activates the expression of potassium transporter operon) and OpuCA (a component of the osmolyte uptake system) (46, 47). Either *gdpP* mutation or *dacA* overexpression leads to c-di-AMP accumulation which negatively influences potassium and other osmolytes uptake; decreased potassium uptake further leads to cell shrinkage, decreased turgor pressure and a smaller cell size (13, 14, 22, 48, 49). A recent study showed that the levels of potassium and free amino acids correlate with antibiotic sensitivity in *Lactococcus lactis* (30). It is hypothesized that an increased c-di-AMP level reduces cellular turgor pressure, generating resistance to osmotic stress and β-lactam antibiotics (30, 45). However, *S. aureus* Δ*gdpP* did not show resistance towards other cell wall-targeting antibiotics, such as moenomycin and bacitracin (22), suggesting that the resistance mechanism may not be limited to the turgor pressure. Another possible explanation is that elevated c-di-AMP alters cell wall structure by increasing cell wall cross-linking and thickness (13, 22, 23). However, the results are inconsistent in the literature in terms of cell wall cross-linking: Δ*gdpP* showed higher cross-linking in a previous study but no change in PG profile as reported in a more recent study (13, 23). Furthermore, c-di-AMP homeostasis has been connected to PG precursor synthesis (32, 50, 51); however, the mechanisms between PG precursor synthesis and β-lactam resistance remain underexplored.

In addition to conferring antibiotic resistance, mutation of *gdpP* can suppress the lethality of *S. aureus* Δ*ltaS* deficient in lipoteichoic acid (LTA) production (13). LTA is a major cell envelope component of Gram-positive bacteria and loss of LTA leads to severe cell growth retardation, cell division defect and cell envelope stress (52, 53). In our previous study, we investigated the role of Δ*ltaS* and Δ*ltaS*Δ*gdpP* in regulating cell wall anchored YSIRK/G-S surface proteins (54). Surface proteins are a group of proteins that are covalently attached to cell wall PG (55). The precursors of surface proteins are secreted across the membrane through the Sec translocon and covalently attached to PG precursor lipid II by the membrane-bound transpeptidase sortase A (SrtA) (56–59). Lipid II-linked surface protein precursors are incorporated into mature peptidoglycan via PBP/SEDS-mediated transpeptidation and transglycosylation during cell wall synthesis (56). Specifically, surface proteins carrying a YSIRK/G-S signal peptide (YSIRK+ proteins) are enriched at the cell division septum and anchored to the septal peptidoglycan (cross-wall) (54, 60). Thus, the dynamic localization of YSIRK+ proteins is coupled with cell cycle and cell wall biosynthesis. We found that the septal trafficking of staphylococcal surface protein A (SpA), an archetype of YSIRK/G-S proteins, was disrupted in strain ANG1786 (Δ*ltaS*Δ*gdpP*) (54). Interestingly, complementation with *ltaS* only restored SpA septal localization, but not its cross-wall localization (data not shown), suggesting that *gdpP* mutation alone may play a role in regulating SpA cross-wall targeting.

In this study, we investigated the role of *gdpP* single mutant in SpA localization, cell wall synthesis and β-lactam resistance. We found that *gdpP* mutation disrupted SpA cross-wall deposition and triggered aberrant foci formation of all four PBPs in *S. aureus*. The mis-localization of PBPs resulted in mis-localized cell wall biosynthesis and correlated with β-lactam antibiotic resistance. Furthermore, Δ*gdpP* exhibited a strong cell cycle retardation leading to significant accumulation of non-dividing cells. We propose that the mis-localization of PBPs and cell cycle arrest contribute to SpA mis-localization and β-lactam antibiotic resistance. Our results not only reveal new mechanisms supporting the cross-wall targeting of YSIRK+ proteins but also provide novel insights into *gdpP*-mediated β-lactam resistance. The mechanisms may be applicable to other bacteria that share conserved regulation by c-di-AMP.

## Results

### The cross-wall deposition of SpA was abolished in Δ*gdpP*

To examine the subcellular localization of SpA, we used two immunofluorescence (IF) methods: cross-wall IF and membrane IF (61–63). In a cross-wall IF experiment, the pre-existing SpA on the bacterial cell wall is removed by trypsin treatment; trypsinized cells are incubated in fresh medium for 20 minutes equal to one round of cell division, allowing the deposition of newly synthesized SpA on the cell surface. Thus, cross-wall IF reveals the localization of surface exposed and cell wall anchored SpA. SpA localizes at the cross-wall of wild-type (WT) cells **(Fig. 1A)**. The membrane IF experiment examines the septal membrane localization of SpA beneath the peptidoglycan layer as trypsinized cells are further treated with cell wall hydrolase lysostaphin, which removes the peptidoglycan layer and separates the two dividing daughter cells, enabling SpA antibody to access the division septum. SpA localizes at the septal membrane in WT cells **(Fig. 1C)**.

**Fig. 1.**
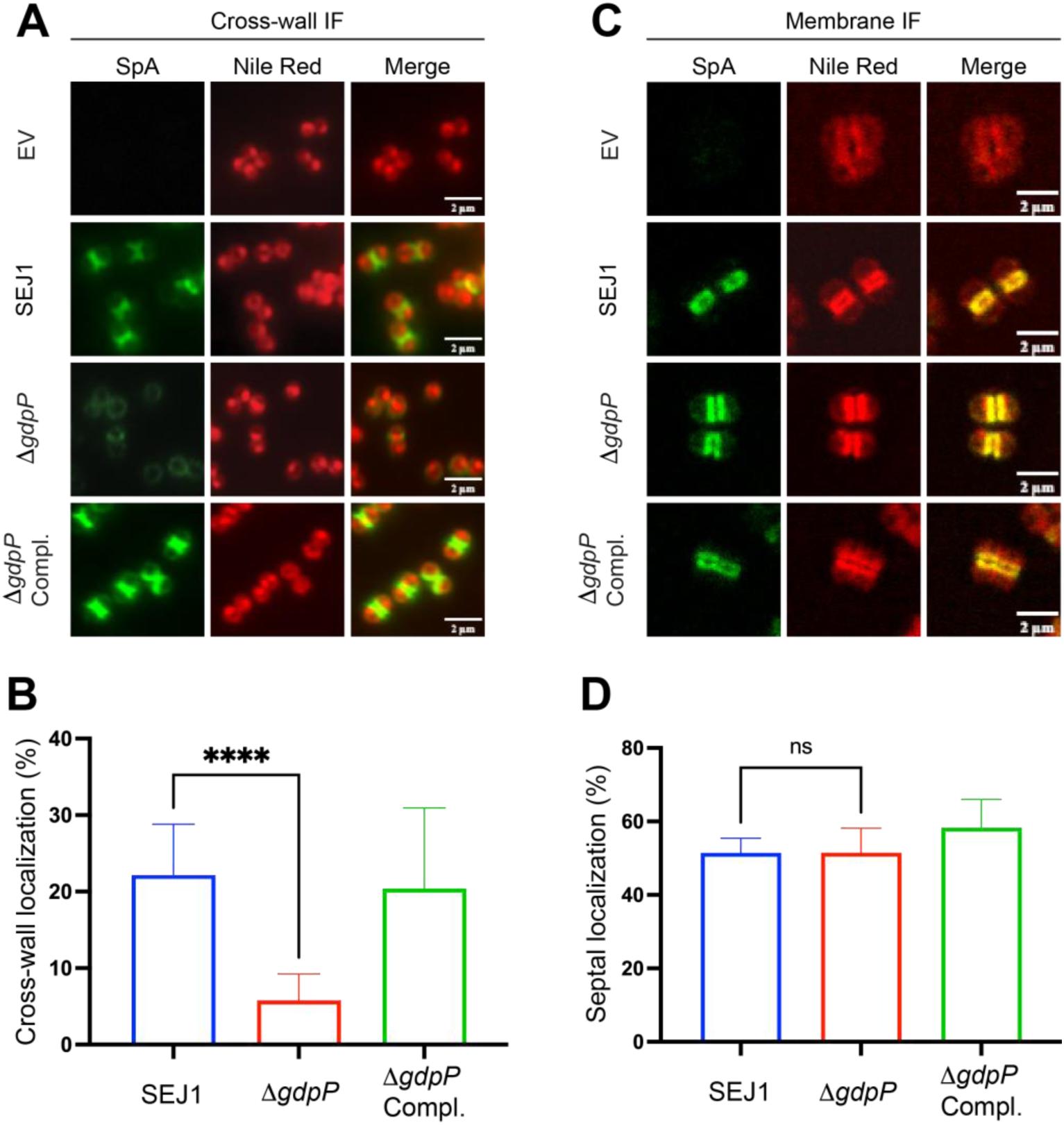
Δ*gdpP* disrupts SpA cross-wall deposition but not its septal membrane localization. (A) Immunofluorescence microscope images showing SpA cross-wall localization in *S. aureus* SEJ1 WT, Δ*gdpP* and *gdpp* complementation strains. SpA expression is controlled by ATc-induction in pCL*itet*-*spa*. EV, empty vector of pCL*itet*. Nile red stained cytoplasmic membrane. Scale bar, 2 µm. (B) Quantification of SpA cross-wall localization from panel A. (C) Immunofluorescence images showing SpA septal membrane localization. (D) Quantification of SpA septal localization. Unpaired t-test with Welch’s correction was performed for statistical analysis: \*\*\*\**p*<0.0001. Representative images and quantification are from three independent experiments.

As previously noted, SpA cross-wall localization remained defective in the Δ*ltaS*Δ*gdpP* strain despite *ltaS* complementation. To further investigate this, we generated a Δ*gdpP::erm* internal deletion mutant **(Fig. S1)**. The *gdpP* gene is located adjacent to two essential genes in a four-gene operon. To avoid potential polar effect, the *erm* marker was removed via the cre-lox system (64). We initially generated the *gdpP* mutant in *S. aureus* strain RN4220 and then transduced Δ*gdpP* to SEJ1, a *spa* deletion mutant of RN4220. The expression of *spa* is induced from a plasmid pCL*itet* with an anhydrotetracycline (ATc)-inducible promoter (54). Cross-wall IF revealed that SpA localized at the cross-wall in SEJ1 but mis-localized all over the cell surface in Δ*gdpP* **(Fig. 1AB)**. SpA fluorescent signal intensity was also reduced in Δ*gdpP*. Complementation by plasmid-borne *gdpP* expression restored the cross-wall localization of SpA, suggesting that the mis-localization of SpA was due to *gdpP* mutation. The septal localization of SpA underneath the peptidoglycan layer was not altered in Δ*gdpP,* as revealed by membrane IF and quantification **(Fig. 1CD)**. These results confirmed that Δ*gdpP* affected SpA localization independent of LtaS. Unlike LtaS that affects membrane localization of SpA (54), deletion of *gdpP* altered the distribution of surface exposed and cell wall anchored SpA.

### Δ*gdpP* displayed aberrant Bocillin-FL foci formation

Once anchored to cell wall PG, the localization of SpA is coupled to cell wall homeostasis, which involves dynamic and balanced synthesis and degradation. Since Δ*gdpP* is resistant to β-lactam antibiotics that target PBPs, we prioritized our research to investigate PBPs-mediated PG synthesis. We first performed Bocillin-FL staining to examine the localization of PBPs in the *gdpP* mutant. Bocillin-FL is a fluorescent penicillin that binds to the transpeptidase domain of all PBPs (65). Fluorescence microscopy showed that Bocillin-FL signals were enriched at the septum in dividing cells and distributed along the cell membrane in non-dividing cells in SEJ1 WT **(Fig. 2A)**. Strikingly, SEJ1Δ*gdpP* exhibited distinct and strong Bocillin-FL foci formation. The signals of Bocillin-FL accumulated as a punctate at one site of the membrane (close to future septum) in non-dividing cells **(Fig. 2A, yellow arrows)**. The Bocillin-FL foci were also found in dividing cells, where the signals of Bocillin-FL distributed as a line at the septa and bright spots were found at the edge or in the middle of the septal line **(Fig. 2A, pink arrows)**. Quantification results revealed that about 65-70% of Δ*gdpP* cells displayed Bocillin-FL foci. Interestingly, almost in all cases, each cell contained only one Bocillin-FL punctate, suggesting that one or more PBPs were mis-localized and aggregated to one single domain in each cell **(Fig. 2B)**. To examine if the phenotype is conserved in *S. aureus*, we transduced Δ*gdpP::erm* to MSSA strains Newman and HG001 and MRSA strain USA300 JE2. Introducing Δ*gdpP* to both MSSA and MRSA strains resulted in smaller cell size and growth defects like SEJ1Δ*gdpP,* confirming the loss of *gdpP* **(Fig. 2)**. The aberrant Bocillin-FL foci formation was observed in all *gdpP* mutants and can be complemented by plasmid-borne *gdpP* expression, indicating that *gdpP* mutation led to mis-localization of PBPs in both MSSA or MRSA.

**Fig. 2.**
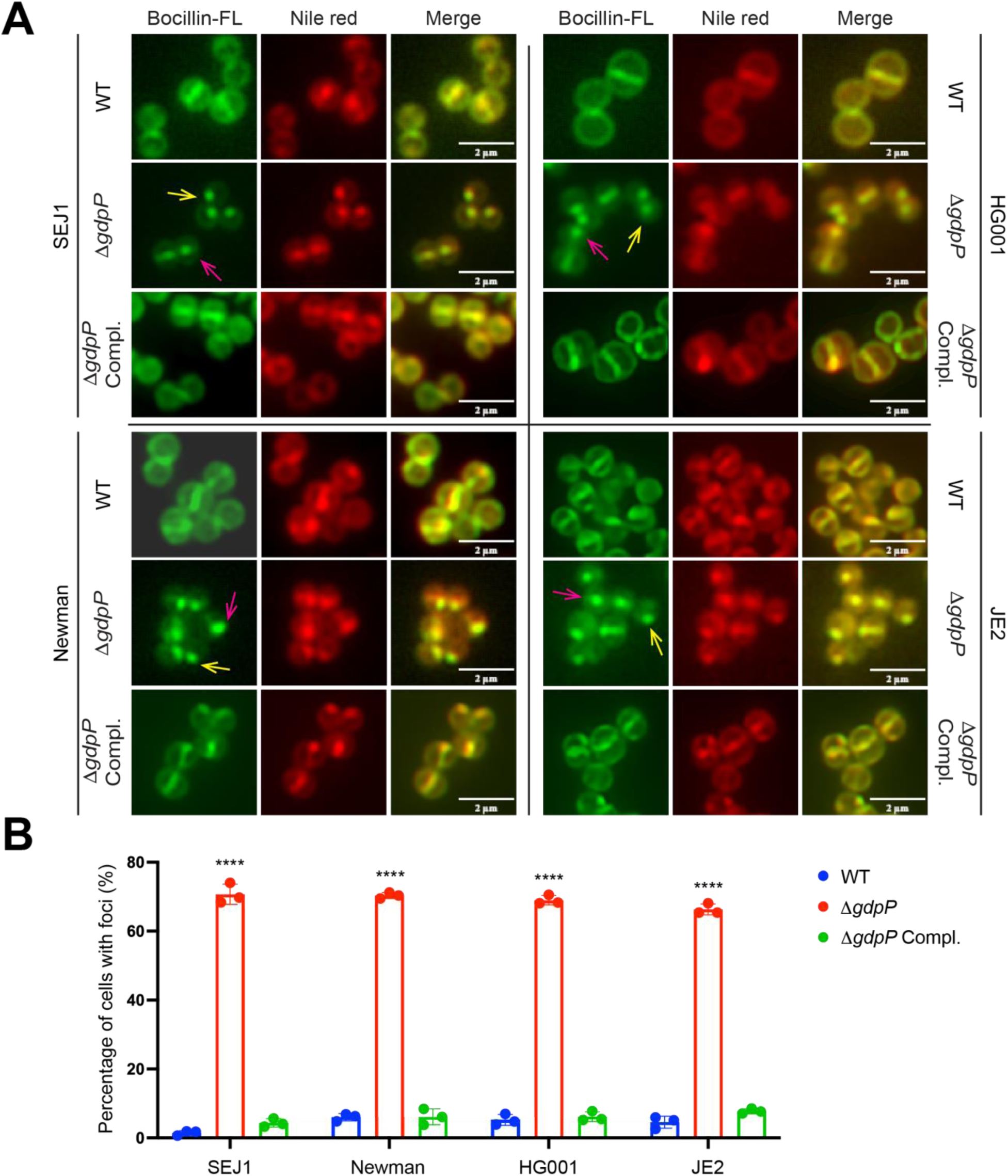
Aberrant Bocillin-FL foci formation in MSSA and MRSA Δ*gdpP*. (A) Fluorescence microscope images of Bocillin-FL staining of WT, Δ*gdpP* and its complementation in different *S. aureus* strains: SEJ1, Newman, HG001, USA300 JE2. SEJ1, Newman and HG001 are MSSA and USA300 JE2 is a MRSA. Nile red stained cytoplasmic membrane. Scale bar, 2 µm. (B) Quantification of foci formation in cells from panel A. Unpaired t-test with Welch’s correction was performed for statistical analysis: \*\*\*\**p*<0.0001. Representative images and quantification are from three independent experiments.

### Overexpression of *dacA* also led to aberrant Bocillin-FL foci formation

We then wanted to study whether the Bocillin-FL foci formation was due to the mutation of *gdpP*, or due to the accumulation of the c-di-AMP. To test this, we overexpressed the c-di-AMP synthetase gene *dacA* in SEJ1, Newman and JE2 via a xylose-inducible promoter from a multicopy plasmid. As a control, glucose was added to repress the expression of *dacA*. Overexpression of *dacA* resulted in smaller cell size like Δ*gdpP*, indicating that the overexpression triggered accumulation of c-di-AMP like the *gdpP* mutant **(Fig. 3)**. In all the strains tested, *dacA* overexpression resulted in significant increase of Bocillin-FL foci formation. These results suggested that the accumulation of c-di-AMP, but not the *gdpP* mutation *per se*, led to Bocillin-FL foci formation.

**Fig. 3.**
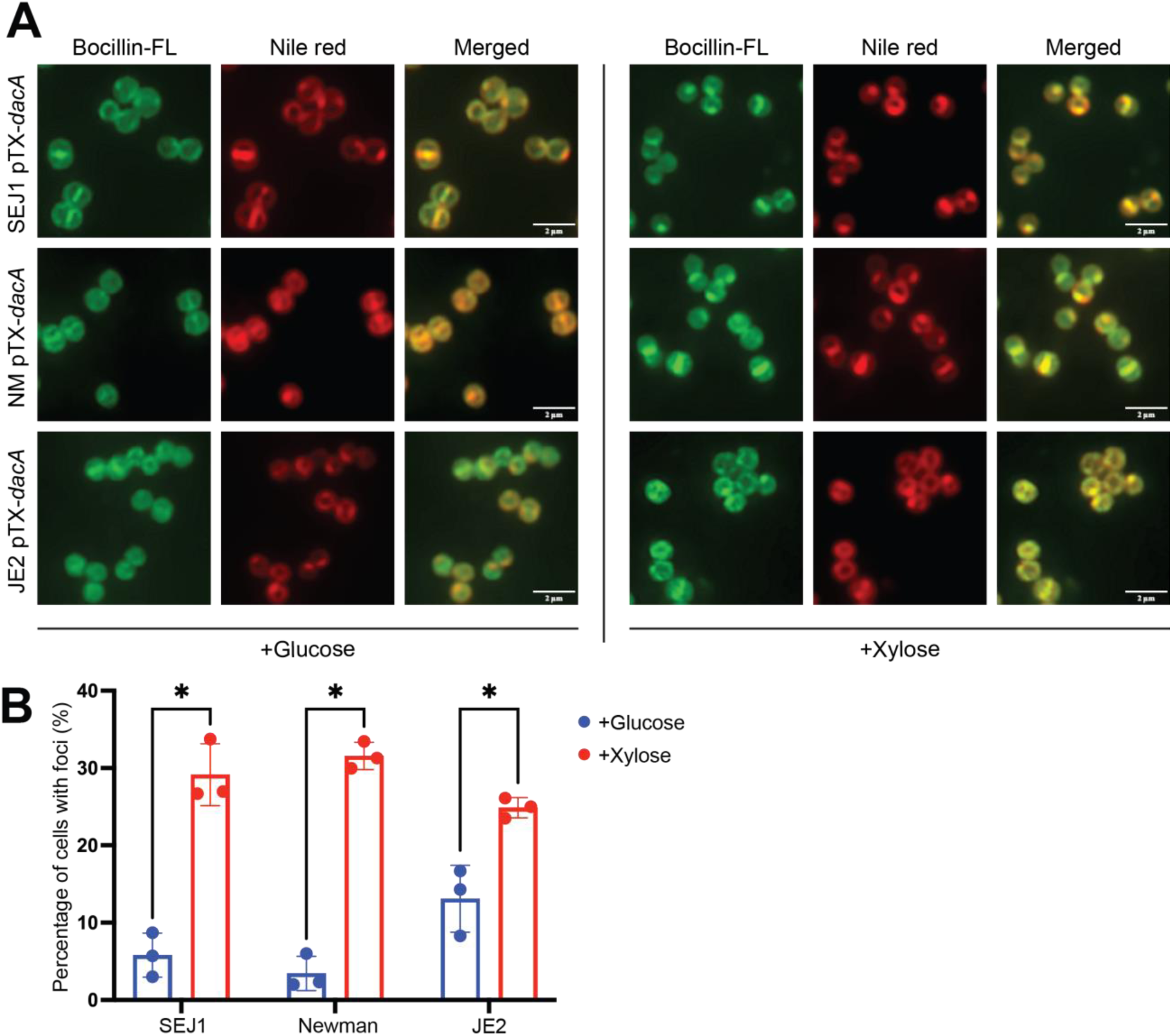
Overexpression of *dacA* led to aberrant Bocillin-FL foci formation. (A) *dacA* overexpressed (+xylose) and repressed (+glucose) cells were stained with Bocillin-FL. Nile red stained cytoplasmic membrane. Scale bar, 2 µm. (B) Quantification of foci formation from panel (A). Unpaired t-test with Welch’s correction was performed for statistical analysis: \**p*<0.05. Representative images and quantification are from three independent experiments.

### All four PBPs are mis-localized in Δ*gdpP*

The genome of *S. aureus* encodes four PBPs, namely PBP1, PBP2, PBP3, and PBP4 (66–68). To further investigate which PBPs were mis-localized in Δ*gdpP*, PBP1-4 were individually tagged with a superfolder green fluorescent protein (sfGFP) and expressed from ATc-inducible promoter from the chromosome. Consistent with previous studies (66–68), all four PBPs localized at the septa of dividing cells and distributed over the cell periphery in non-dividing cells in SEJ1 WT. Intriguingly, all four PBPs formed distinct foci in Δ*gdpP* **(Fig. 4A)**. Similar to the results of Bocillin-FL staining, the foci of PBPs were deposited as a single punctate at one site of the membrane in each cell. Nearly 40% of cells displayed sfGFP-PBP2 foci, which is the highest among the four PBPs. SfGFP-PBP1, sfGFP-PBP3, and PBP4-sfGFP showed lower percentage (about 25%) of foci formation in *ΔgdpP*, but all were significantly higher compared to 2-5% of foci formation in WT **(Fig. 4B)**. This suggested that the essential cell wall synthase PBP2 may be mostly affected in *ΔgdpP*. The percentage of PBPs foci formation is lower than that of Bocillin-FL foci, which could be due to the higher stability and sensitivity of Bocillin-FL staining compared to GFP-tagging. Complementation of *gdpP* restored the localization of PBPs comparable to the WT, confirming the role of GdpP in modulating proper PBPs localization.

**Fig. 4.**
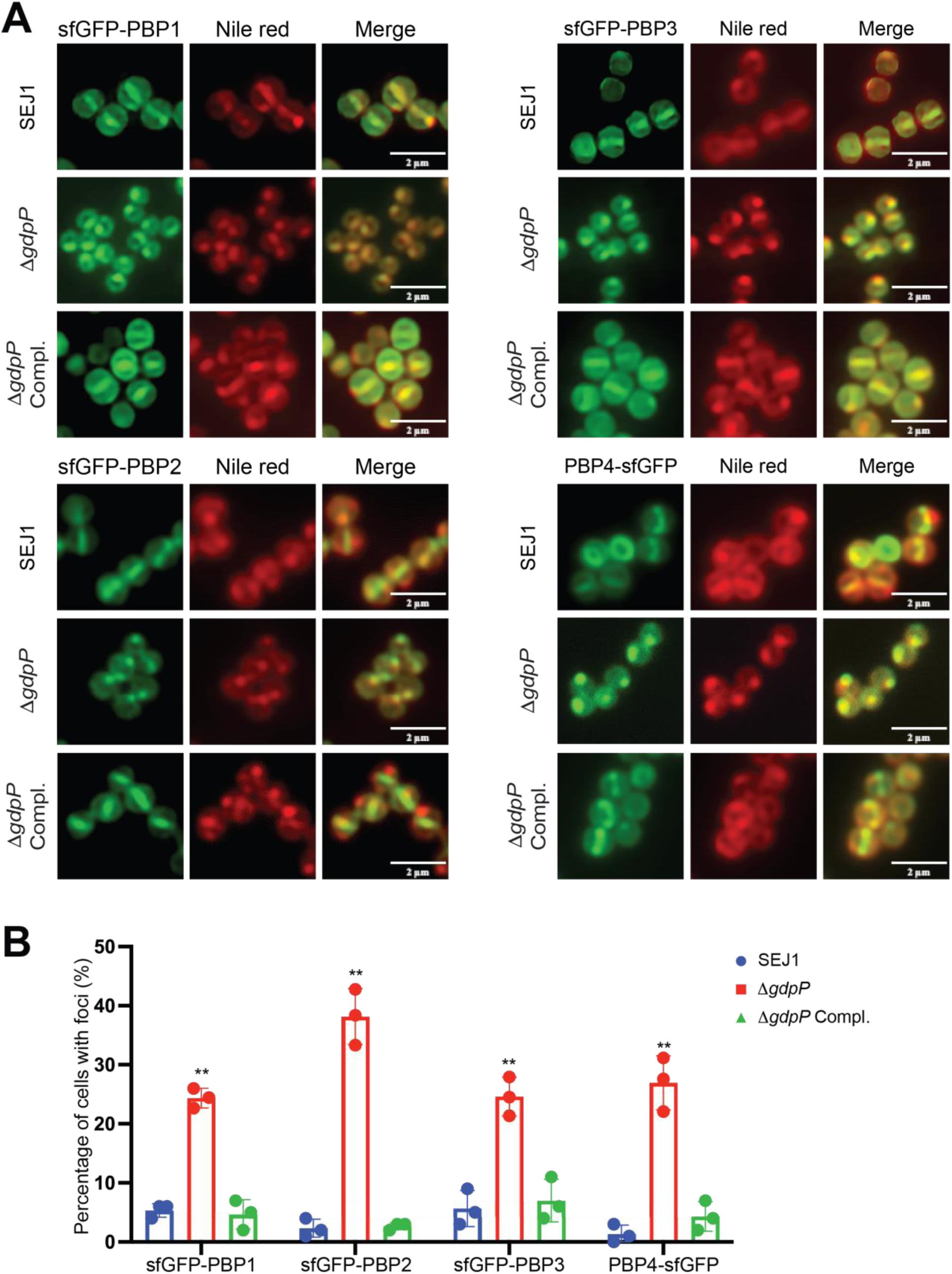
Aberrant foci formation of all four native PBPs (PBP1-4) in *ΔgdpP*. (A) Fluorescent microscope images of sfGFP-PBP1, sfGFP-PBP2, sfGFP-PBP3, and PBP4-sfGFP in SEJ1, SEJ1 Δ*gdpP*, and SEJ1 Δ*gdpP* complement strains. Nile red stained cytoplasmic membrane. Scale bar, 2 µm. (B). Quantification of foci formation in cells from (A). Unpaired t-test with Welch’s correction was performed for statistical analysis: \*\**p*<0.005; \*\*\**p*<0.0005. Representative images and quantification are from three independent experiments.

### Mis-localization of PBPs led to mis-localized cell wall synthesis

The aberrant localization of SpA and PBPs prompted us to further investigate the localization of cell wall synthesis. To test this, we sequentially labeled the *S. aureus* cells with three fluorescent D-amino acid (FDAAs): HADA (blue), RADA (red), OGDA (green) (60). The FDAAs are actively incorporated into the cell wall via the transpeptidase activity of PBPs and the method has been widely used to label active cell wall synthesis sites (60, 69–71). In our experiment, log-phase staphylococcal cultures were incubated with each FDAA for 20 min, equivalent to roughly one round of cell division. Sequential FDAA staining reveals cell wall synthesis dynamics across two rounds of cell division. In WT *S. aureus* cells, HADA stains the old cell wall of the two daughter cells (half-moon shape), RADA stains the new cell wall synthesis (Y’ or ‘V’ shape) and OGDA stains the next round of septal cell wall synthesis in two daughter cells (straight lines) **(Fig. 5)**. In comparison to WT, the three FDAAs often co-localized and RADA and OGDA frequently formed foci at the cell periphery of Δ*gdpP* cells **(Fig. 5, arrows)**, indicating delayed and mis-localized cell wall synthesis. The mis-localized FDAA incorporation is likely due to the mis-localization of PBPs, which was observed in all strain backgrounds and quantification revealed a statistically significant difference **(Fig. 5B)**.

**Fig. 5.**
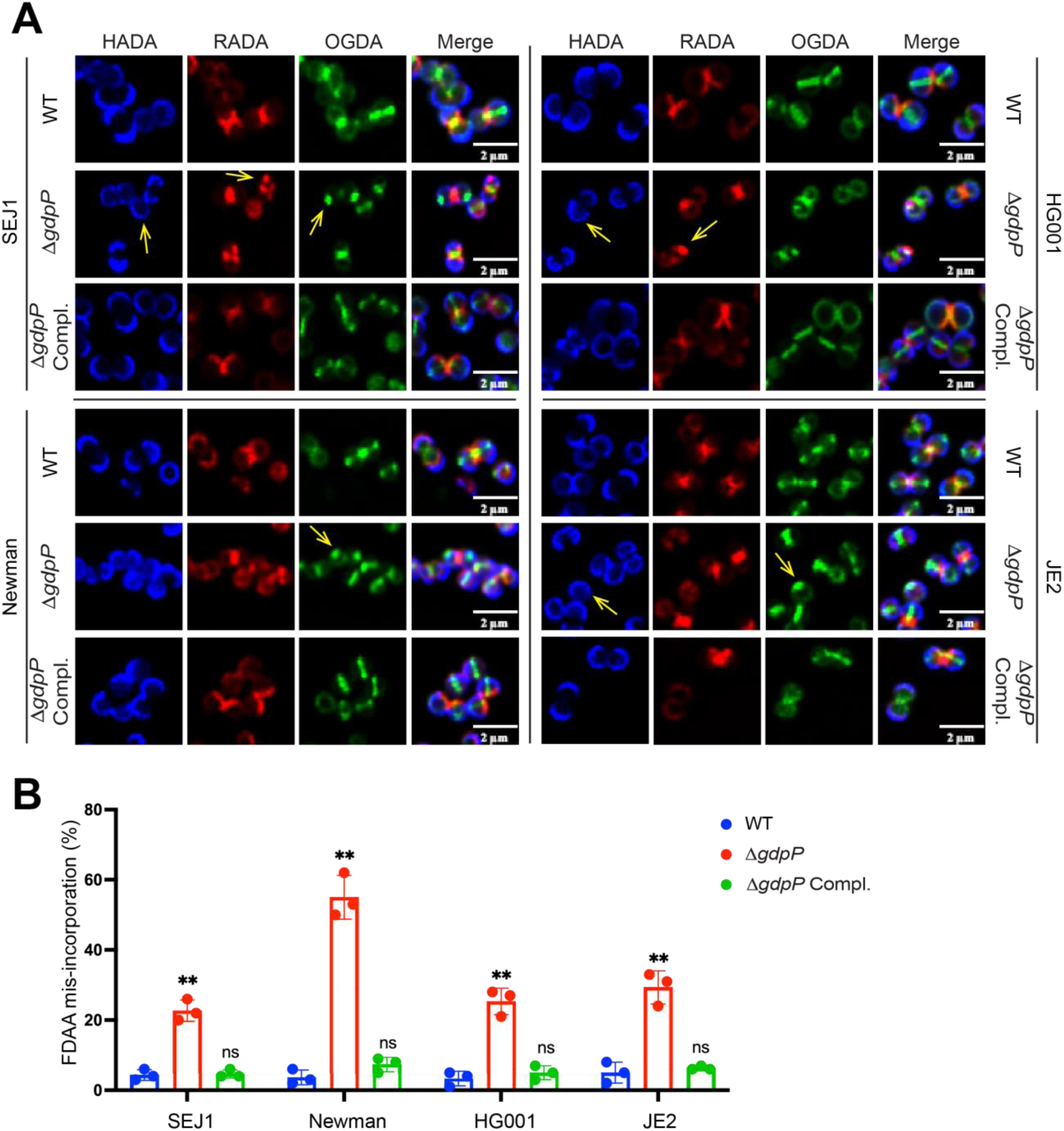
The dynamic localization of cell wall biosynthesis was altered in Δ*gdpP*. (A) WT, Δ*gdpP*, and *gdpP* complementation cells were sequentially incubated with three FDAAs: HADA (blue), RADA (red) and OGDA (green) and imaged by fluorescence microscopy. Scale bar, 2 µm. (B) Quantification of FDAA mis-incorporation. An unpaired t-test with Welch’s correction was performed for statistical analysis: \*\**p*<0.005. Representative images and quantification are from three independent experiments.

We then examined whether the mis-localization of PBPs would affect the overall cell wall PG structure. PG analysis from previous studies showed inconsistent results: *ΔgdpP* was reported to have increased PG cross-linking in an earlier study while a recent study showed no difference (13, 23). Since PBP4 is the primary contributor for highly cross-linked PG in *S. aureus* (72, 73), we included Δ*pbp4* and generated a Δ*gdpP*Δ*pbp4* double mutant. The cell wall peptidoglycan from SEJ1 WT, Δ*gdpP*, Δ*pbp4* and Δ*gdpP*Δ*pbp4* were purified, digested with mutanolysin (N-acetylmuramidase) and analyzed by high-performance liquid chromatography (HPLC). Δ*gdpP* showed a slight increase in PG cross-linking, but quantification and statistical analysis revealed no significant differences **(Fig. 6)**. As expected, Δ*pbp4* exhibited a significant decrease in PG cross-linking. There was a small increase in PG cross-linking in Δ*gdpP*Δ*pbp4* compared to Δ*pbp4,* but the increase was not statistically significant. Overall, the data obtained suggested that Δ*gdpP* did not significantly alter PG structure and cross-linking.

**Fig. 6.**
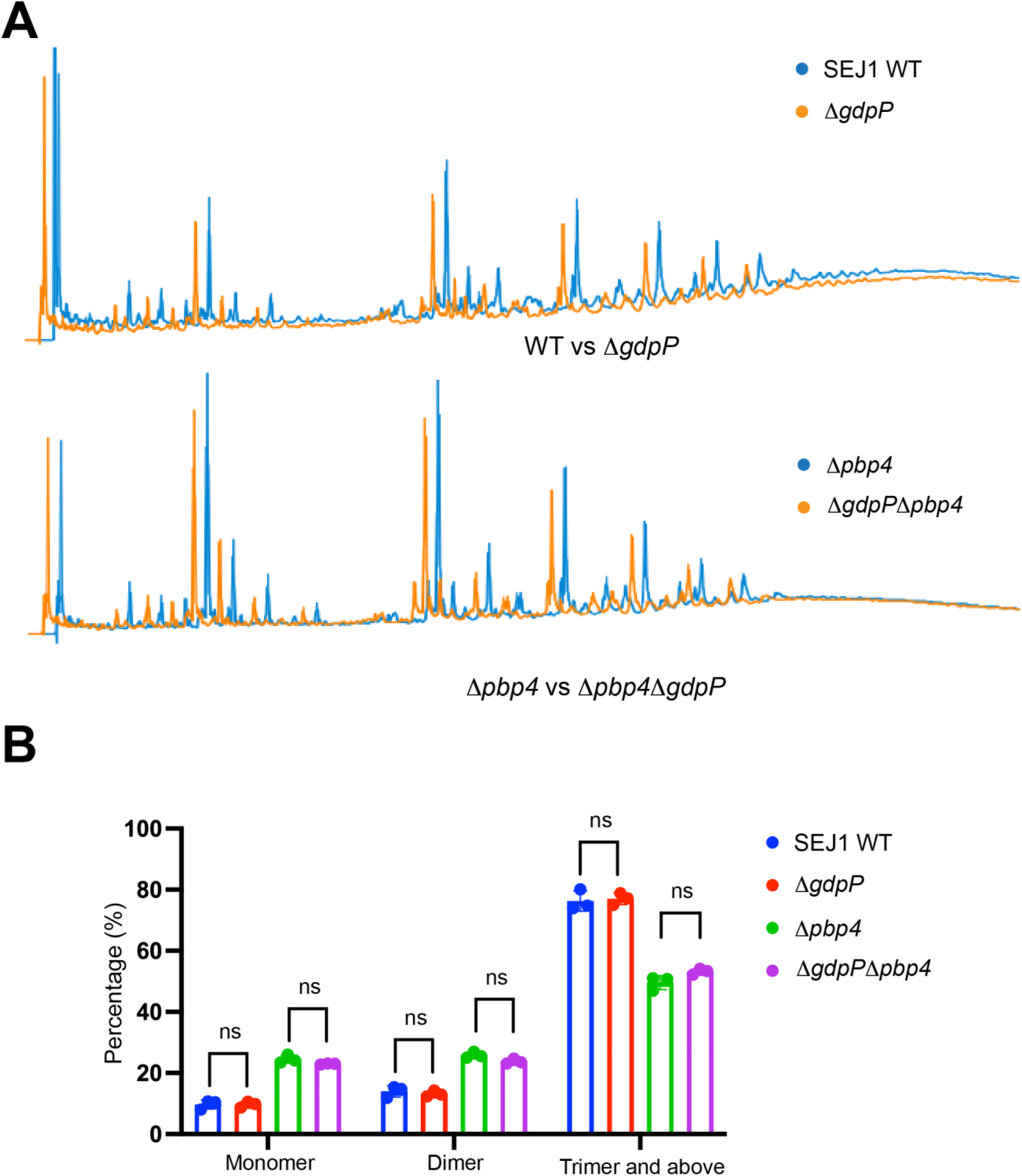
Cell wall cross-linking was not altered in Δ*gdpP*. **(A)** HPLC analysis of cell wall cross-linking in SEJ1 WT, Δ*gdpP*, Δ*pbp4*, and Δ*gdpP*Δ*pbp4*. **(B)** Quantification of the area under the curve of monomers, dimers, and trimers and above from (A). Unpaired t-test with Welch’s correction was performed for statistical analysis.

### Δ*gdpP* is significantly retarded in cell division

From our microscope experiments, we noticed that most of the Δ*gdpP* cells were not dividing. To further investigate it, we performed cell cycle analysis. The cell cycle of staphylococcal cells has been defined as three phases (60, 74): the phase 1 cells exhibit no septa, which have completed their previous cell division and not started the next cycle; the phase 2 cells exhibit incomplete septa, which have just initiated cell division; the phase 3 cells have completed septa and exhibit an elongated shape. Compared to the WT, Δ*gdpP* showed 15-20% increase in phase 1 cell population, and 5% and 10-15% decrease in phase 2 and 3 cell populations. The alteration was observed in all strain backgrounds and was statistically significant **(Fig. 7)**. The accumulation of phase 1 non-dividing cells strongly indicates a defect in cell division initiation. To further analyze cell division initiation, we examined ZapA-sfGFP localization **(Fig. 8)**. ZapA is an early cell division protein that promotes FtsZ polymerization and divisome formation (53, 75, 76). The localization of ZapA indicates the sites of divisome formation at the early stage of cell division. Microscopy and quantification analysis revealed that the frequency of ZapA septal localization (ZapA ring formation) was decreased by 7% in Δ*gdpP* compared to WT, indicating a defect in cell division initiation. However, the decrease of ZapA septal ring formation (7%) is not as dramatic as the decrease of dividing cells (15-20%) **(Fig. 7)**, suggesting that additional mechanisms may contribute to cell division defect of Δ*gdpP*.

**Fig. 7.**
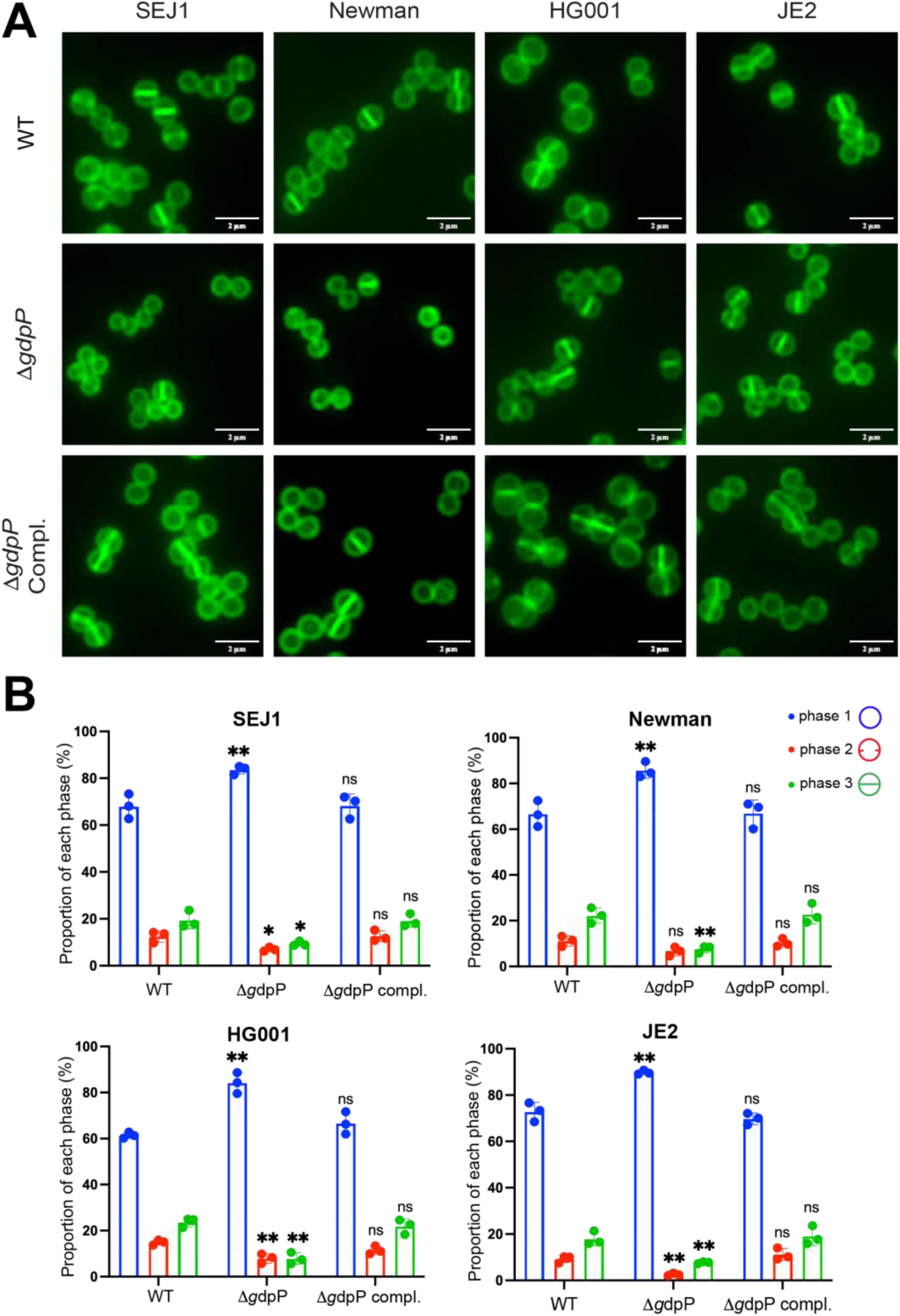
Δ*gdpP* exhibits strong cell cycle retardation. (A) Van-FL labelling of WT, Δ*gdpP*, and *gdpP* complementation in *S. aureus* SEJ1, Newman, HG001, and JE2. Scale bar, 2 µm. (B) Quantification of cells from different stages of the cell cycle: with no septum (P1), a partial septum (P2), or a complete septum (P3). Asterisks on top of each sample indicate statistical analysis result between WT and the sample. Unpaired t-test with Welch’s correction was performed for statistical analysis: \**p*<0.05; \*\**p*<0.005. Representative images and quantification are from three independent experiments.

**Fig. 8.**
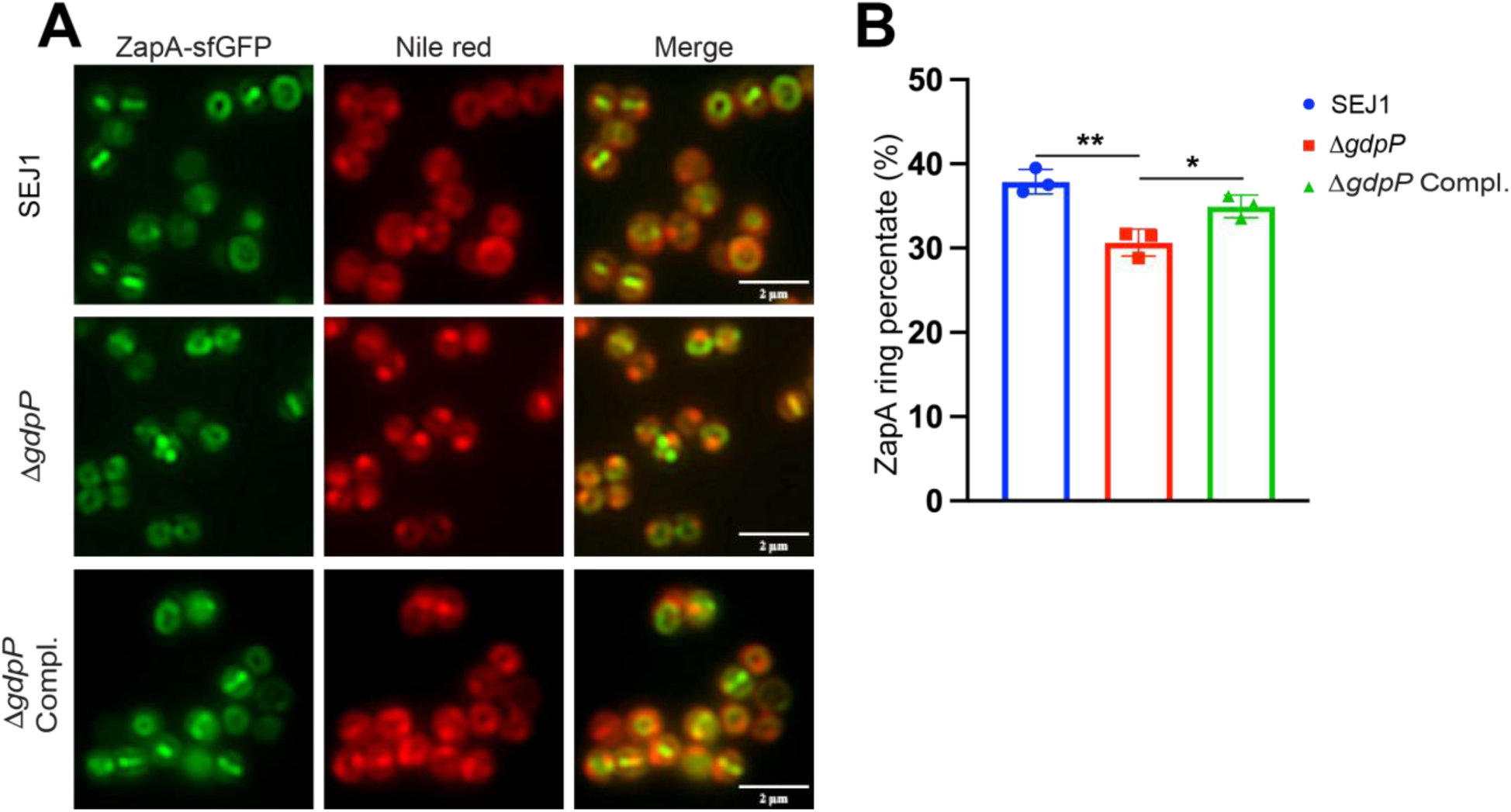
Decreased ZapA septal ring formation in Δ*gdpP*. (A) The localization of ZapA-sfGFP in SEJ1 WT, Δ*gdpP*, and *gdpP* complementation. Nile red stained cytoplasmic membrane. Scale bar, 2 µm. (B) Quantification of ZapA septal ring formation. Unpaired t-test with Welch’s correction was performed for statistical analysis: \**p*<0.05; \*\**p*<0.005. Representative images and quantification are from three independent experiments.

### The mis-localization of PBPs correlates with β-lactam resistance

Finally, we aimed to examine if the mis-localization of PBPs contributed to β-lactam resistance. We first tested the MIC of Δ*gdpP* towards β-lactam antibiotics (penicillin G and oxacillin) and vancomycin **(Fig. S2)**. Both β-lactam antibiotics and vancomycin inhibit PG cross-linking albeit with different mechanisms: β-lactam antibiotics target the transpeptidase domain of PBPs whereas vancomycin binds to d-alanyl-d-alanine terminus of the muropeptides. Consistent with previous study (13), Δ*gdpP* was resistant towards β-lactam antibiotics penicillin and oxacillin, but not vancomycin **(Fig. S2)**. MRSA JE2Δ*gdpP* exhibited higher resistance towards penicillin (MIC 8 µg/ml) compared to MSSA SEJ1Δ*gdpP* (MIC 0.47 µg/ml). We then treated both SEJ1 and JE2 WT and Δ*gdpP* cells with various concentrations of penicillin and stained with Bocillin-FL. Treatment with lower concentrations of penicillin (0.01 µg/ml for SEJ1 and 0.05 µg/ml for JE2) in the WT cells readily triggered enlarged cell size and aberrant division septa formation **(Fig. 9)**. These morphological changes indicated the killing activity of penicillin in weakening the cell wall and inducing cell lysis. In comparison, low concentration of penicillin did not alter the morphology of Δ*gdpP* nor the Bocillin-FL foci formation. Increasing penicillin concentration gradually triggered the morphological change and the killing of Δ*gdpP*. Importantly, the Bocillin-FL foci formation gradually decreased with increasing concentrations of penicillin **(Fig. 9B)**. At high concentration of penicillin close to the MIC (0.04 µg/ml for SEJ1Δ*gdpP* and 5 µg/ml for JE2Δ*gdpP*), the Δ*gdpP* cells were significantly enlarged with irregular septa and the Bocillin-FL foci completely disappeared, indicating that the high MIC concentration of penicillin disrupted the Bocillin-FL foci and triggered killing. Thus, the presence of Bocillin-FL foci correlated with the sensitivity/resistance of Δ*gdpP* to penicillin.

**Fig. 9.**
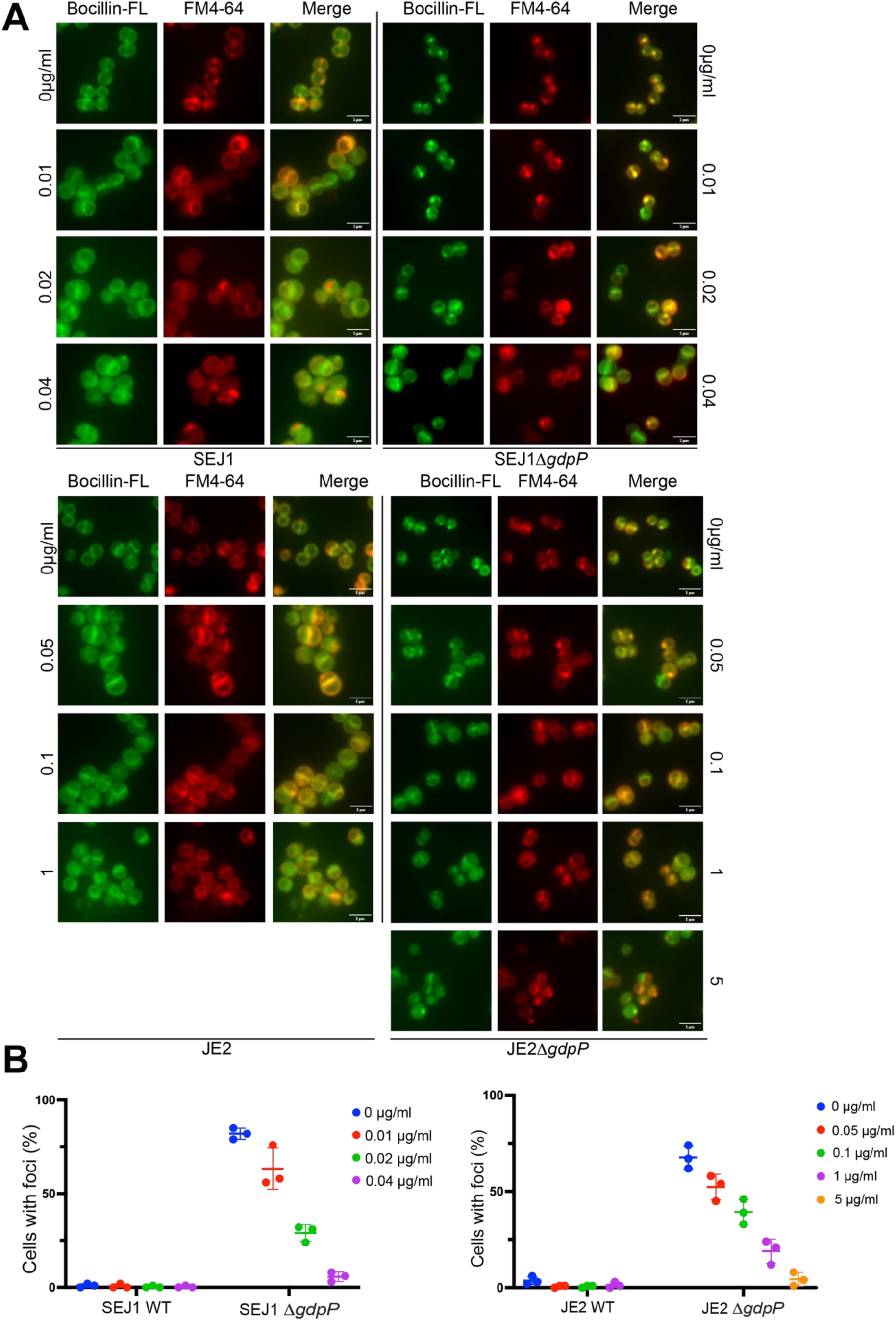
The foci of PBPs correlate with β-lactam resistance. (A) Bocillin-FL staining of SEJ1(MSSA) and JE2(MRSA) WT, Δ*gdpP*, and *gdpP* complementation strains treated with different concentrations of penicillin. FM4-64 stained cytoplasmic membrane. Scale bar, 2 µm. (B) Quantification of Bocillin-FL foci formation.

Taken together, we showed here that *S. aureus* Δ*gdpP* disrupted YSIRK+ protein SpA cross-wall deposition and triggered aberrant foci formation of all four PBPs. The mis-localization of PBPs resulted in mis-localized cell wall synthesis, but did not significantly alter the overall PG structure and cross-linking. Concomitantly, Δ*gdpP* was resistant to PBP-targeting β-lactam antibiotics, but not to cross-linking targeting antibiotics such as vancomycin. The foci formation of PBPs correlated with the killing activity of β-lactam antibiotic. Δ*gdpP* also exhibited a strong defect in cell division initiation with an accumulation of non-dividing cells. We concluded that the mis-localization of PBPs and cell division arrest of Δ*gdpP* dysregulated SpA cross-wall trafficking and rendered resistance to β-lactam antibiotics **(Fig. 10)**.

**Fig. 10.**
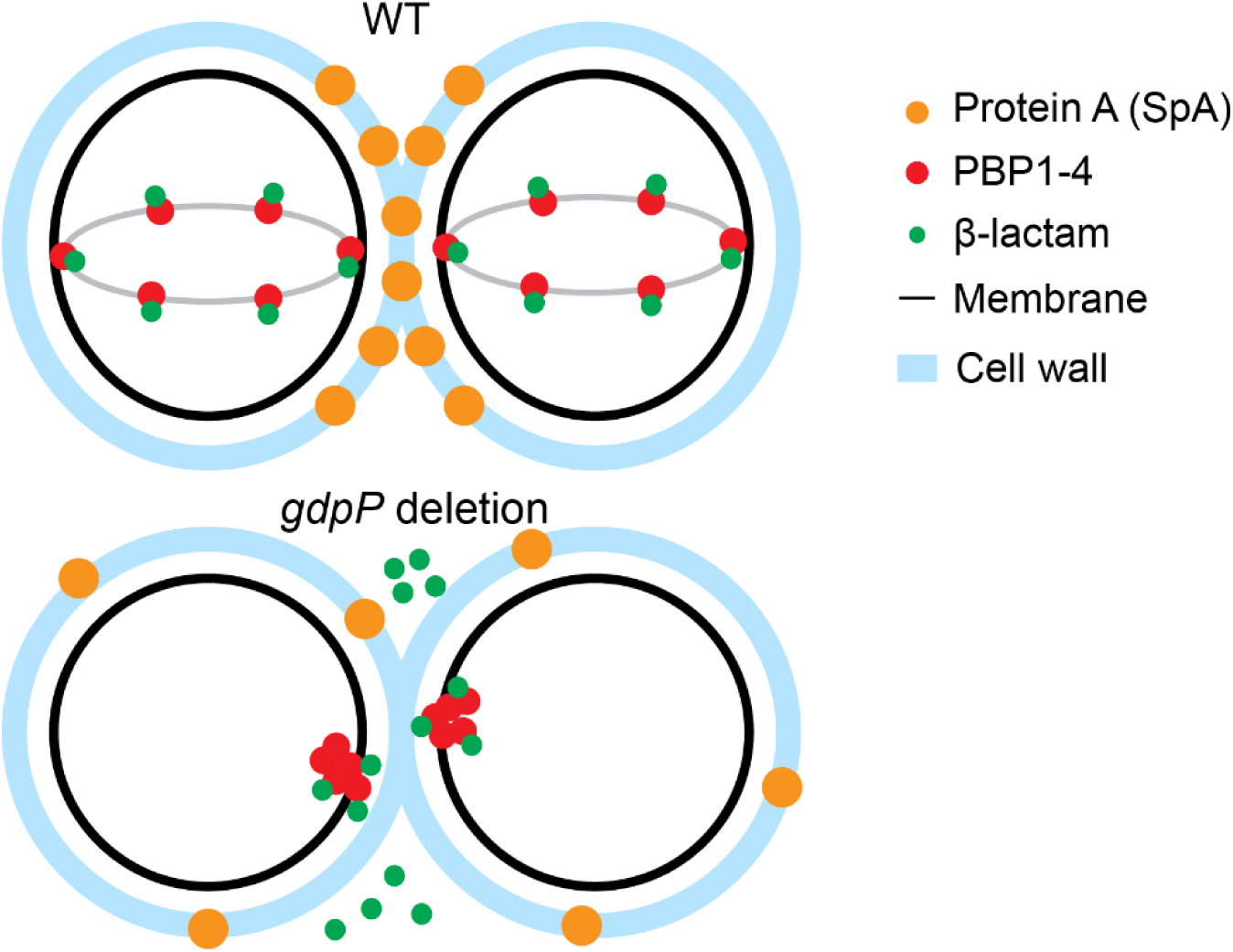
Model of how Δ*gdpP* regulates PBPs localization, YSIRK+ protein SpA trafficking, and β-lactam resistance. In WT cells, PBP1-4 are localized at the septum in dividing cells and distributed along the cell membrane in non-dividing cells. In Δ*gdpP* cells, the cell division is retarded and PBPs accumulate as distinct single foci. The mis-localization of PBPs and reduction in septation decreases the binding of β-lactam antibiotics to septal localized PBPs, making β-lactam antibiotics less effective. Furthermore, the mis-localized PBPs are unable to incorporate SpA precursor to cross-wall, resulting in SpA mis-localization.

## Discussion

While cumulative studies have reported the correction of *gdpP* mutation and the level of c-di-AMP with β-lactam resistance, the underlying mechanisms are largely unknown. β-lactam antibiotics target the transpeptidase activity of PBPs essential for PG cross-linking. In *S. aureus*, new PG synthesis mainly occurs at the septum and all the four native PBPs (PBP1-4) of *S. aureus* are known to localize at the septum during PG synthesis (66–68, 77). PBP1 is a high-molecular-weight (HMW) class B PBP with only transpeptidase activity, and it is essential for cell septation and separation (78, 79). PBP2 is a HMW class A PBP and is bifunctional with both transpeptidase and transglycosylase activities (80). It is also essential for cell division and remains localized at division septum upon initiation stage of daughter cells separation (77). PBP3 and PBP4 are non-essential PBPs with only transpeptidase activity (72, 81, 82). Class B PBPs PBP1 and PBP3 form pairs with monofunctional transglycosylase FtsW and RodA (known as SEDS proteins) respectively, which participate in septal and sidewall PG synthesis (83). PBP4 is a class C PBP that is recruited to the division septum after PBP1 and PBP2 (84). It has carboxypeptidase activity and transpeptidase activity that contribute to the generation of highly crosslinked PG (85).

Strikingly, all four PBPs are mis-localized as aberrant foci in Δ*gdpP.* As both Bocillin-FL staining and sfGFP-tagged PBPs displayed one focus per cell, PBP1-4 likely aggregate together and localize at the same position in Δ*gdpP*. Consistent with the previous report (15), there was no significant change in the protein abundance of PBPs when we extracted membrane proteins and stained with Bocillin-FL (data not shown). There was discrepancy in terms of PG cross-linking in the literature (13, 23). We repeated the cell wall PG analysis, and our results are in line with the recent study (23): there was no significant difference in PG cross-linking between WT and Δ*gdpP.* The PG analysis has limitations as it analyzes the cross-linking in a whole cell population, which may not reveal the difference at single cellular level. Nevertheless, the results suggest that the PBPs are largely functional in Δ*gdpP.* Thus, neither the expression nor the activity of PBPs is significantly altered in Δ*gdpP*. Our cell cycle analysis and ZapA-GFP localization results indicate that Δ*gdpP* exhibits defects in cell division initiation leading to accumulation of non-dividing cells and reduction of dividing cells. We propose that the mis-localization of PBPs and the decrease of actively dividing cells contribute to β-lactam resistance in Δ*gdpP*. Mis-localization of PBPs at the cell periphery and the accumulation of non-dividing cells decrease the binding of β-lactam antibiotics to septal localized PBPs, making the β-lactam antibiotics less effective.

We propose that the mis-localization of PBPs in Δ*gdpP* dysregulates SpA cross-wall localization. Surface proteins are important components of staphylococcal cell envelope, which play crucial roles in staphylococcal pathogenesis (86). Surface proteins that carry a YSIRK/G-S signal peptide are known to be enriched at the septal membrane and deposited at the cross-wall (54). However, the detailed mechanisms are unclear. We previously demonstrated that LtaS-mediated LTA synthesis spatially regulates SpA trafficking, as revealed by both membrane IF and cross-wall IF (54, 61). Here we show that Δ*gdpP*, the suppressor of LtaS, regulates SpA cross-wall deposition, but not its septal membrane localization. Thus, the mechanism of GdpP-mediated regulation differs from that of LtaS. Interestingly, we observed a similar phenotype in the mutant lacking D-alanylation of teichoic acids: the *dlt* mutant abolishes SpA cross-wall localization but does not affect SpA septal membrane localization (54). However, Δ*dlt* displays distinctly different cell morphology compared to Δ*gdpP*: the *dlt* mutant cells are enlarged with increased cell lysis and decreased cross-linking (54). The results suggest that both Δ*gdpP* and Δ*dlt* regulate the later step of SpA biogenesis, but the mechanisms may differ. Most likely, Δ*gdpP* dysregulates SpA cross-wall deposition via mis-localization of PBPs while Δ*dlt* alters cell lysis and the PG structure. However, we cannot rule out that other mechanisms, such as increased autolysis in Δ*gdpP* (13, 51), may play a role in regulating SpA localization.

What remains unknown is how PBPs are mis-localized in the *gdpP* mutant. The results that PBPs were mis-localized upon *gdpP* deletion and *dacA* overexpression indicated that elevated c-di-AMP level dysregulates PBPs localization. Although we have not directly measured the level of intracellular c-di-AMP, previous studies have provided sufficient evidence that the mutation of *gdpP* or overexpression of *dacA* results in c-di-AMP accumulation (22, 48). Decreased cell size (Fig. 5A) and reduced growth rate (data not shown) observed in our strains further support the effect of c-di-AMP accumulation. Various mechanisms have been shown to regulate the localization of PBPs in *S. aureus,* including the availability of transpeptidase substrates, wall teichoic acids, and divisome components (66, 77, 84, 87). We propose that the mis-localization of PBPs in Δ*gdpP* is an indirect consequence of c-di-AMP accumulation. In *S. aureus*, c-di-AMP directly binds to proteins responsible for potassium and compatible solutes transportation (46, 47). Accumulation of c-di-AMP reduces intracellular potassium and other osmolytes, which decreases cellular turgor pressure and leads to cell shrinkage (30, 45). The cell shrinkage may alter the fluidity and the structure of cytoplasmic membrane, which in turn leads to aggregation of membrane proteins. Since all four PBPs are mis-localized, the mechanism might be more generic, potentially affecting the localization of other cell division-related proteins. The precise mechanism underlying PBP mis-localization in *ΔgdpP* remains to be further investigated.

## Materials and methods

### Bacterial strains and growth conditions

Strains and plasmids used in this study are listed in Table S1. *E. coli* strain DC10B, DH5α and DH5α λpir were grown in Luria–Bertani broth (LB) or on LB agar plate supplemented with the appropriate antibiotics. 100 µg/ml of ampicillin (Amp) and 10 µg/ml trimethoprim (Tmp) was used for plasmid selection in *E. coli*. *S. aureus* strains were grown in tryptic soy broth (TSB) or on tryptic soy agar plate (TSA) supplemented with the appropriate antibiotics: 7 µg/ml of chloramphenicol (Chl), 10 µg/ml of erythromycin (Ery), 10 µg/ml of trimethoprim (Tmp), and 25 µg/ml of tetracycline (Tet). 200 ng/ml of ATc was used to induce gene expression from the *P_tet_* promoter in pCL*itet*. 1% glucose and xylose were used to repress and induce *P_xyl_* promoter in pTX plasmid. If not specified, overnight cultures were grown at 37°C with appropriate antibiotics and 1:100 refreshed to TSB the next day. Appropriate inducers were added to the refreshed cultures. Refreshed cultures were grown for 2–3 hours to mid-log phase (OD_600_ of 0.8) and proceeded with different experiments.

### Construction of plasmids and strains

To construct Δ*gdpP*, an internal deletion of 1529 bp of the *gdpP* gene (*SAOUHSC_00015*) including its SD sequence was deleted. The gene knockout vector pKOR1 plus cre-lox marker removal system was used to generate the *gdpP* mutant (64, 88). To generate pKOR1-*gdpP::erm-lox,* primers 1143/1144 were used to amply the upstream DNA fragment, primers 1147/1148 were used to amplify *lox66-ermB-lox71* from *smc::erm-lox* (89), and primers 1145/1146 were used to amplify downstream DNA fragment. The three fragments were digested by XhoI&PstI, ligated and amplified by primers 1143&1148. The PCR product (up+erm+down) was digested by EcoRI&EcoRV and ligated to pKOR1. The resulting knockout plasmid pKOR1-*gdpP::erm-lox* was confirmed by restriction enzyme digestion and sequencing. The homologous recombination process followed the previous protocol plus *erm^+^* counter selection (88). The final deletion mutant of *gdpP* was confirmed by PCR and sequencing. SEJ1Δ*gdpP*, NewmanΔ*gdpP*, HG001Δ*gdpP*, and JE2Δ*gdpP* were constructed by transducing Δ*gdpP::erm* to WTstrains. SEJ1Δ*gdpP* pCL*itet-spa* was constructed by transducing Δ*gdpP::erm* to SEJ1 pCL*itet-spa* (54). To build SEJ1Δ*gdpP*::-, pRAB1 was transformed to SEJ1Δ*gdpP::erm*. The *erm* cassette and subsequently the pRAB1 plasmid were removed according to the procedures described in previous studies (64). The resulting SEJ1Δ*gdpP*::-was confirmed by PCR and sequencing. SEJ1Δ*pbp4* and SEJ1Δ*gdpP*Δ*pbp4* were generated by phage transduction of Δ*pbp4::Tn* from Nebraska Transposon Mutant Library (NTML) (90).

To construct GFP-tagged PBP1-4, primers 702/648 were used to PCR amplify *pbp1* from SEJ1 genomic DNA. The PCR product was ligated with NheI/BglII digested pCL*itet*-*sfgfp-sfAgfp* (60) via Gibson assembly (NEB), generating pCL*itet-sfgfp-pbp1*. Primers 704/650 were used to PCR amplify *pbp2* from SEJ1 genomic DNA. The PCR product was restricted by NheI/BamHI and ligated with plasmid pCL*itet*-*sfgfp-sfgfp* restricted by NheI/BglII, generating pCL*itet-sfgfp-pbp2*. Primers 705/652 were used to PCR amplify *pbp3* from SEJ1 genomic DNA. The PCR product was restricted by NheI/BamHI and ligated with plasmid pCL*itet*-*sfgfp-sfgfp* restricted by NheI/BglII, generating pCL*itet-sfgfp-pbp3*. The construction of pCL*itet-pbp4-sfgfp* has been described in our previous study (60). The *gdpP* complementation plasmid was generated as follows: primers 626/627 and 628/629 were utilized to amplify the *gdpP* operon promoter (290 bp upstream of SAOUHSC_00014) and the *gdpP* gene from SEJ1 genomic DNA. The PCR product of *gdpP* promoter and *gdpP* gene were digested with NheI/SacI and SacI/BamHI respectively, and the pKK30 plasmid were digested with NheI/BamHI. Digested products were ligated to generate pKK30-P*gdpP-gdpP*, simplified as pKK30*-gdpP*. For *dacA* overexpression, primers 1149/1150 were used to amplify *dacA* from RN4220 genomic DNA. The PCR product was digested with BamHI/SacI and ligated into the pTX plasmid carrying a xylose-induced promoter to generate pTX-*dacA*. To generate pALFS1-*zapA-sfgfp*, the *zapA-sfgfp* fusion was cut out from pJB67-*zapA-sfgfp* (75) by SalI/BamHI digestion ligated with pALFS1, a modified pJB67 with the antibiotic *erm^R^*cassette replaced by *chl^R^.* The positive clones confirmed by DNA sequencing were electroporated to the desired *S. aureus* strains. Primers used in this study are listed in Table S2.

### Immunofluorescence microscopy and quantification

SpA immunofluorescence microscopy was performed based on the protocols described in previous studies (54, 61–63). Briefly, in the cross-wall IF experiment, mid-log phase cultures were collected and treated with trypsin. Trypsinized cells were washed and incubated in fresh medium containing trypsin inhibitor for 20 min at 37°C to allow SpA regeneration. Subsequently cells were fixed and stained with SpA primary antibodies and fluorescently labeled secondary antibodies (anti-rabbit IgG conjugated to Alexa Fluor 488) (Invitrogen). In a membrane IF experiment, trypsinized cells were fixed and digested with lysostaphin (AMBI) on the glass slides for 2 min. The samples were fixed again in methanol and acetone followed by immunostaining. All samples were stained in the dark with Hoechst 33342 DNA dye (Invitrogen) and Nile red (Sigma). Fluorescent images were captured by Nikon Scanning Confocal Microscope Eclipse Ti2-E with HC PL APO 63×oil objective (1.4 NA, WD 0.14 mm).

Quantification of SpA localization: at least 300 cells for each sample from three independent experiments were analyzed with ImageJ (91). The total cell numbers with SpA signals and cells displaying cross-wall/septal SpA localization were counted. Unpaired t-test with Welch’s correction was performed for statistical analysis using GraphPad Prism: *p <0.05; **p <0.005; ***p <0.0005; ****p <0.0001.

### *dacA* overexpression

Bacterial cells with pTX-*dacA* were incubated in TSB overnight and subsequently washed twice with basic medium without glucose (10 g/L Soy Peptone, 5 g/L Yeast extract, 5 g/L NaCl, 1 g/L K_2_HPO_4_, pH=7.2) without glucose. Washed cells were inoculated 1:100 in 10 ml basic medium without glucose in a flask and grew for 2 hours to reach OD_600_ of 0.5. 1% xylose was added to induce *dacA* expression, while 1% glucose was added to repress its expression as a control. After an additional two hours of incubation, cells were harvested for further study.

### Fluorescence microscopy of PBPs’ localization

For Bocillin-FL staining, bacterial cells at OD_600_ of 1.5 were harvested by centrifugation. The cell pellets were washed with phosphate buffered saline (PBS) (137 mM NaCl, 2.7 mM KCl, 10 mM Na_2_HP_4_, 1.8 mM KH_2_PO_4_, pH=7.4) and incubated with 10 µM Bocillin-FL penicillin sodium salt (Invitrogen) in TSB at 37°C for 30 min. To study the localization of sfGFP-tagged PBPs, bacteria cells were incubated TSB supplemented with 200 ng/mL ATc for 3 hours to induce the expression of sfGFP-tagged fusion proteins. Subsequently, samples were harvested by centrifugation, washed and resuspended in PBS. The cells were then fixed in 1 ml of 50% fixation solution (2.5% paraformaldehyde and 0.01% glutaraldehyde in PBS), washed twice with PBS, and applied to poly-L-lysine-coated 8-well glass slides (MP Biomedicals). Excess and non-adherent cells were washed away with PBS, and the samples were then strained with Hoechst 33342 DNA dye and Nile red or FM™ 4-64 Dye (Invitroegn). Samples were covered with 15 µl of SlowFade™ diamond antifade mountant (Invitrogen) and sealed with glass coverslips. All images were captured by Nikon Scanning Confocal Microscope ECLIPSE Ti2-E equipped with the HC PL APO 63×oil objective (1.4 NA, WD 0.14 mm).

Quantification and statistical analysis: at least 300 cells for each sample from three independent experiments were analyzed by ImageJ software (91). The total cell numbers and cells displaying Bocillin-FL foci or GFP foci were counted. Unpaired t-test with Welch’s correction was performed for statistical analysis using GraphPad Prism: *p <0.05; **p <0.005; ***p <0.0005; ****p <0.0001.

### Fluorescent D-amino acid (FDAA) staining

FDAA staining was performed as described earlier (60). Bacteria were cultured to mid-log phase and adjusted to OD_600_ of 0.8. Cells were then incubated with 500 µM HADA (TOCRIS) in TSB for 20 min at 37°C with shaking, followed by a single wash using 0.01 M PBS. Subsequently, cells were incubated in TSB containing 500 µM RADA (TOCRIS) for another 20 minutes at 37°C, washed once with PBS, and then incubated in TSB with 500 µM OGDA (TOCRIS) for 20 minutes at 37°C, followed by washing with PBS. Washed cells were fixed with fixation solution for 20 min at room temperature, after which they were washed with PBS. Then, 50 µl of the resuspended cells were applied to poly-L-lysine coated glass slides and allowed to dry at room temperature. Samples were covered with 15 µl of SlowFade™ diamond antifade mountant and sealed with glass coverslips. Images were captured by Nikon Scanning Confocal Microscope ECLIPSE Ti2-E equipped with the HC PL APO 63×oil objective and analyzed in ImageJ (91).

### Van-FL staining and cell cycle quantification

Overnight cultures were refreshed 1:100 in TSB and incubate to OD_600_ of 1.0. Cells were fixed with 1 ml 50% fixation solution and washed twice with PBS and then applied to poly-L-lysine-coated 8-well glass slides. Excess and non-adherent cells were washed away with PBS and samples were then strained with 1 µg/ml BODIPY™ FL Vancomycin (Invitrogen) to stain the cell wall.

Quantification of cell size and cell cycle: at least 300 cells per sample from three independent experiments were analyzed in ImageJ (91). The staphylococcal cell cycle has been previously defined (74): phase 1, cells without a septum; phase 2, cells with a partial septum; phase 3, cells with a complete septum and elongated shape. Unpaired t-test with Welch’s correction was performed for statistical analysis using GraphPad Prism.

### Cell wall purification and HPLC

Muropeptide purification from *S. aureus* was conducted according to established protocols with some modifications (92–94). Bacterial cells were incubated overnight and refreshed in 1 L of TSB with an initial OD_600_ of 0.1. Cells were harvested at OD_600_ of 0.6 and boiled in 5% SDS for 15 minutes to remove non-covalent bound proteins followed by several washes with water to remove SDS. Staphylococcal cells were mechanically lysed by bead beater with 0.1 mm silica beads, beating 3450 RPM at 115V for 30s with 8 cycles (MiniBeadBeater-16, Biospec). Subsequently, silica beads were removed by centrifugation, and cells were incubated with 10 µg/mL DNAse and 50 µg/mL RNAse at 37°C for 2 hours with shaking. An overnight digestion with 50 µg/mL trypsin was performed to remove surface proteins. Then, cells were treated with 48% HF for 48 hours at 4°C to remove WTA, followed by washing several times with water to remove HF. Purified peptidoglycan was resuspended in Na-Phosphate buffer (50 mM Na_2_HPO_4_ and 50 mM NaH_2_PO_4_, adjust pH to 7.0), normalized to an OD_600_ of 3.0, and treated with 50 U/100 µl Mutanolysin (Millipore Sigma) overnight at 37°C to obtain muropeptides. Muropeptides were reduced by sodium borohydride in 500 mM sodium borate (pH=9.0) for 20 minutes and the reaction was quenched by adjusting the pH to 3.0 with 30% phosphoric acid. Then, samples were analyzed by reverse phase HPLC (Agilent 1260 Infinity II) using the InfinityLab Poroshell 120 EC-C18 (4.6 x 150 mm, 2.7 µm) HPLC column. Elution was performed using a gradient from 5% to 30% methanol in 100 mM Na-Phosphate buffer (0.4 ml/min) for 150 minutes and monitored at 205 nm. Quantification of PG profile: three independently grown cultures were utilized to purify the cell wall and analyzed by HPLC. The areas of muropeptide peaks were integrated and quantified using Agilent Data Analysis version 2.7 and presented as a percentage of the total.

### Antibiotic resistance and microscopy

The strip tests were used to examine antibiotic resistance. Overnight bacterial cultures were refreshed 1:100 in TSB and grow to OD_600_ of 1.0. Cells were then normalized to OD_600_ of 0.1 and applied to the TSA plates with and without 2% NaCl with a cotton swab. Penicillin G and vancomycin strips were placed in TSA plates, and oxacillin strips were placed in TSA plates supplemented with 2% NaCl (95). Images of plates were taken after 24 hours incubation. Samples for microscopy were prepared as follows: SEJ1 and SEJ1Δ*gdpP* were treated with 0.01 µg/ml, 0.02 µg/ml, 0.04 µg/ml penicillin G; JE2 and JE2Δ*gdpP* were treated with 0.05 µg/ml, 0.1 µg/ml, 1 µg/ml, and 5 µg/ml penicillin G. After 3 hours incubation, cells were harvested, washed with PBS, and the OD_600_ was normalized to 1.5. The cell pellets were washed with PBS and stained with Bocillin-FL as described above. Fluorescence microscopy images were captured by Nikon Scanning Confocal Microscope ECLIPSE Ti2-E equipped with the HC PL APO 63×oil objective. Images from three independent experiments and at least 300 cells per sample were quantified and analyzed in ImageJ (91).

## Acknowledgements

This work is supported by NIH/NIGMS-R35GM146993. We thank members of the Yu lab for critically reading the manuscript and providing suggestions.

---

**Fig. S1.**
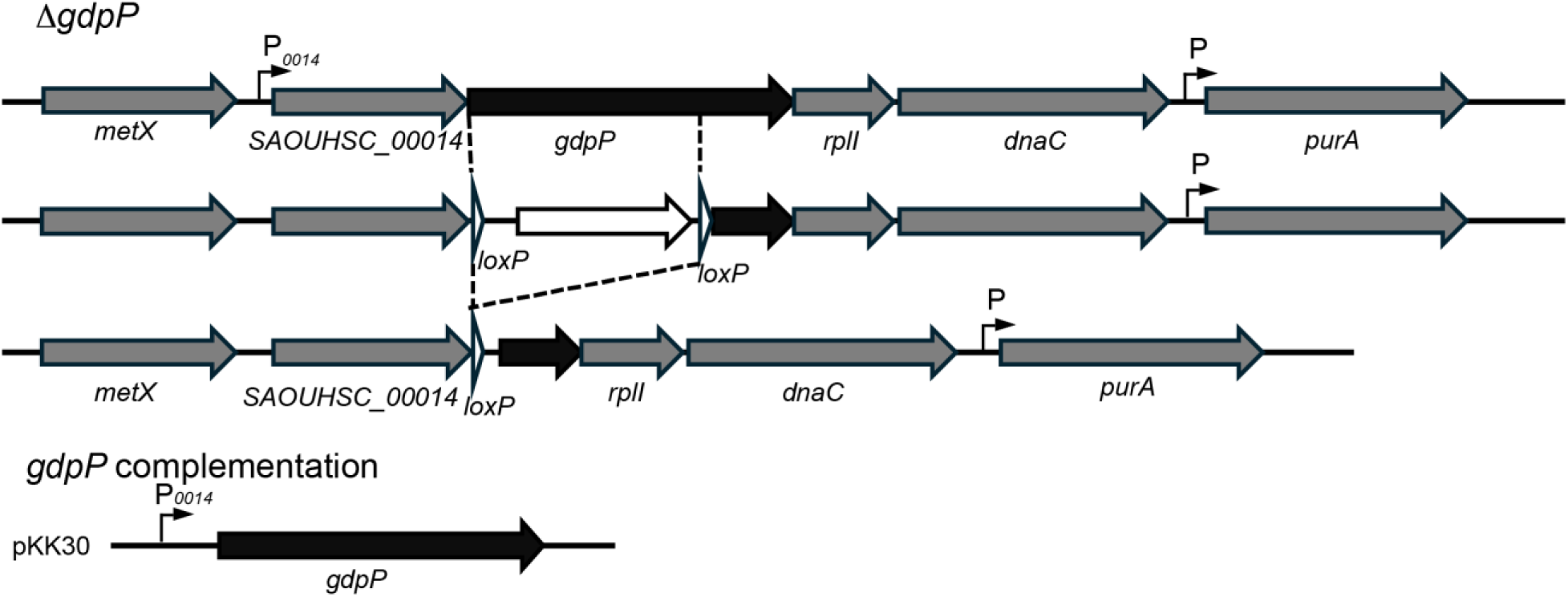
Construction of SEJ1Δ*gdpP*. The *gdpP* coding sequence, except for 449 bp at the 3’ end, was replaced by *ermB* cassette flanked with *lox* sites. *ermB* cassette was further removed by Cre recombinase. pKK30-P*_gdpP_-gdpP* plasmid was used to complement Δ*gdpP*.

**Fig. S2.**
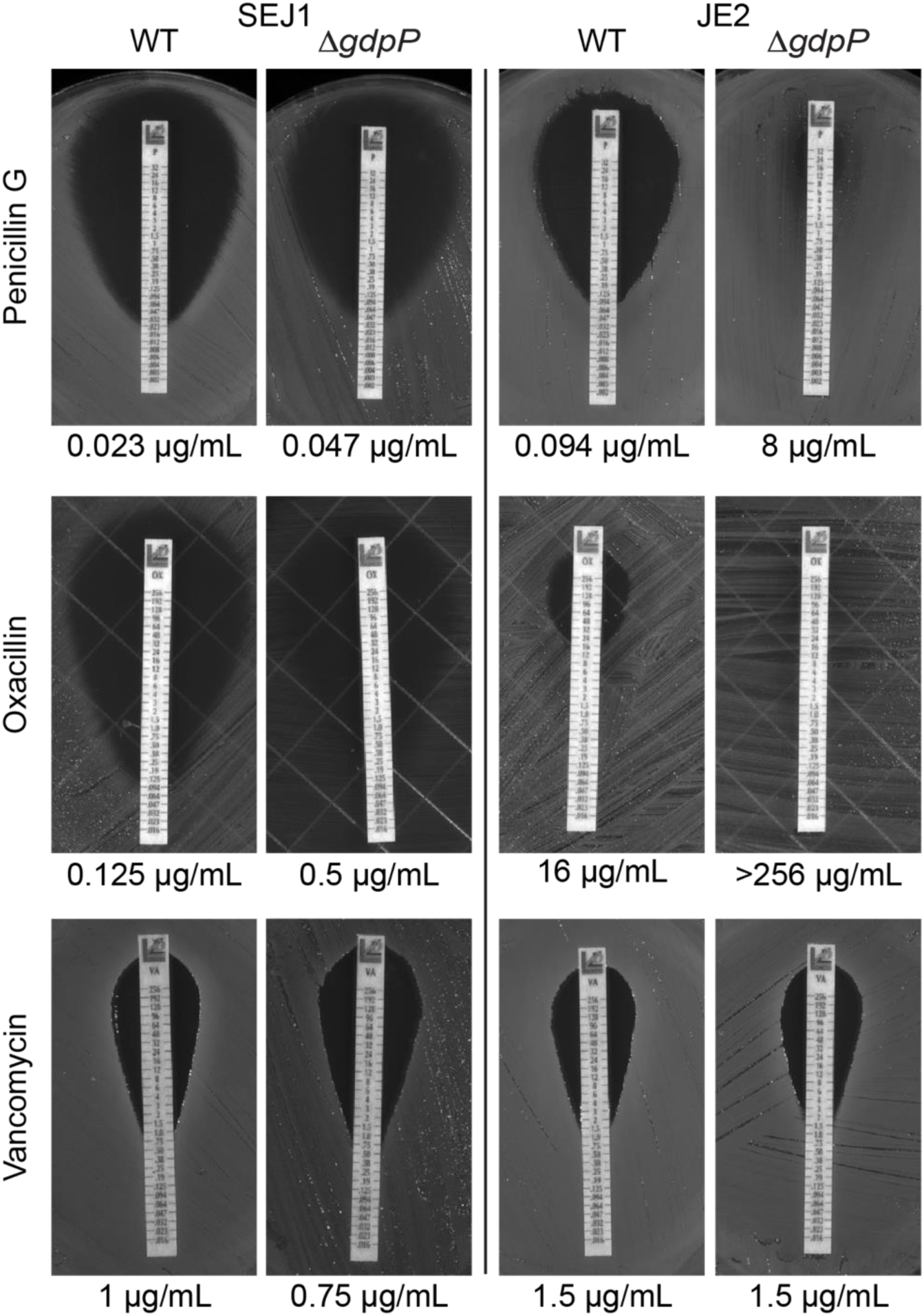
Antibiotic strip test showing an increased penicillin and oxacillin resistance in Δ*gdpP* strains, with no significant change in vancomycin resistance. MIC values was shown below the strip test images.

**Table S1.**
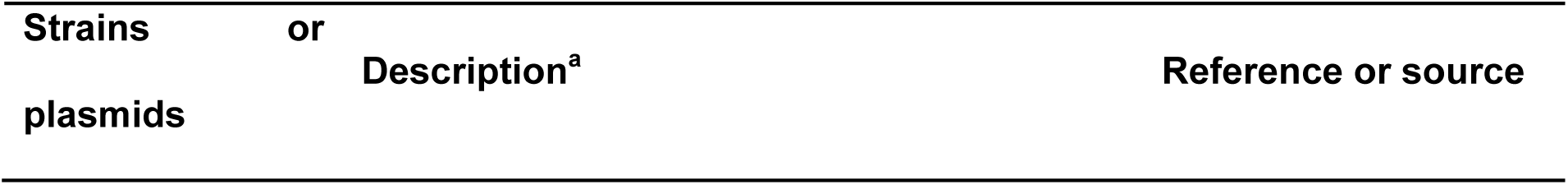
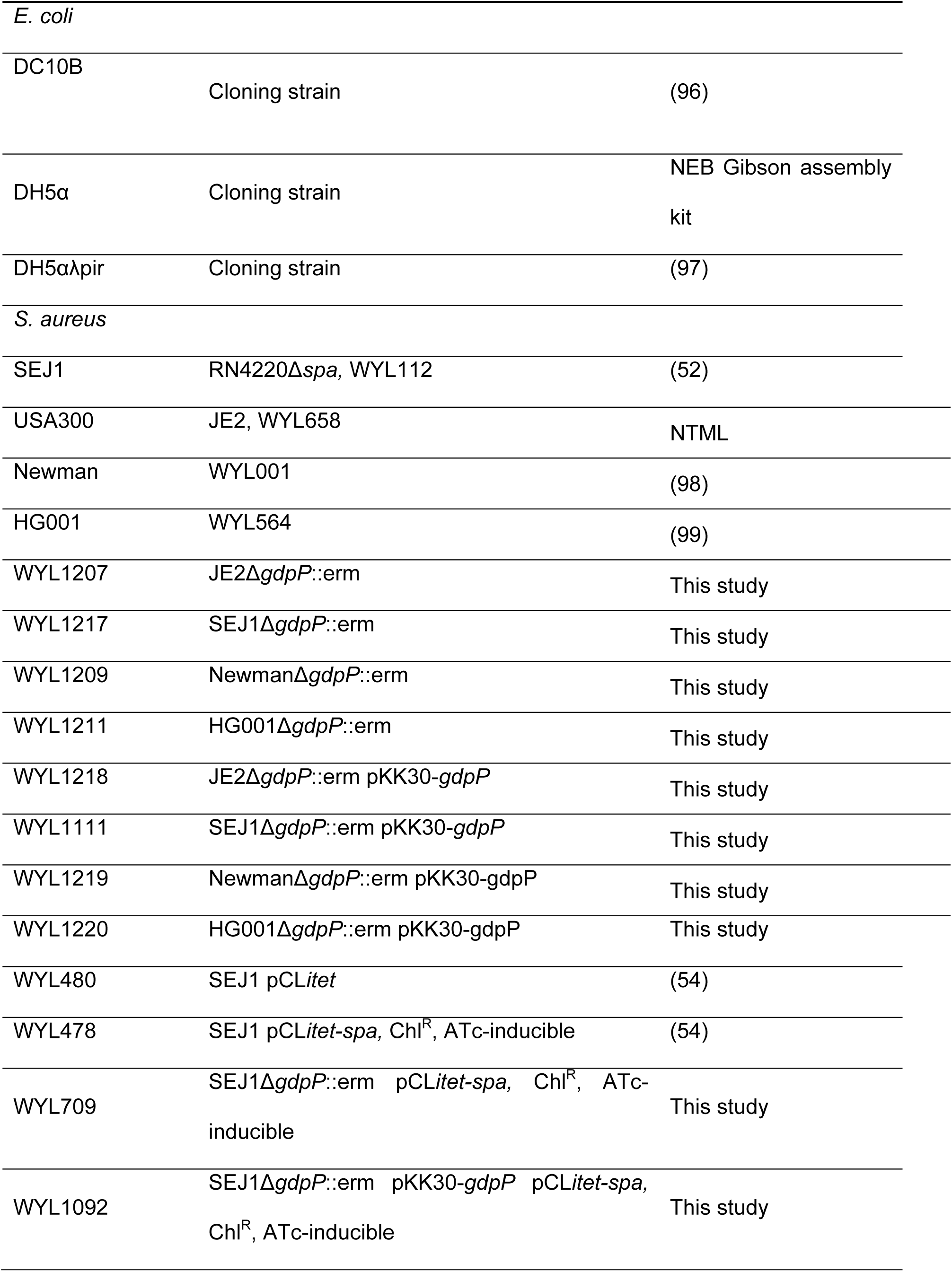
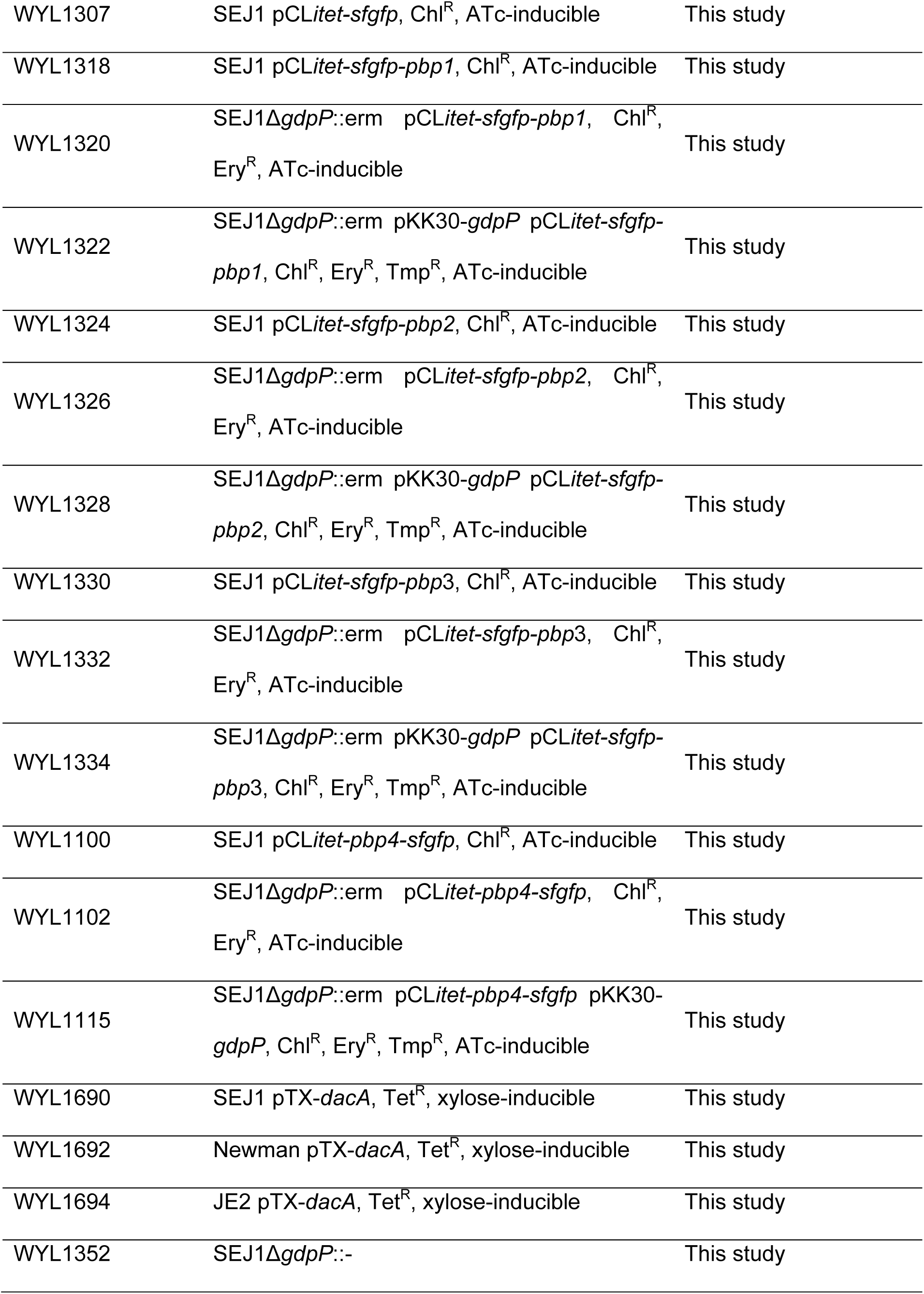
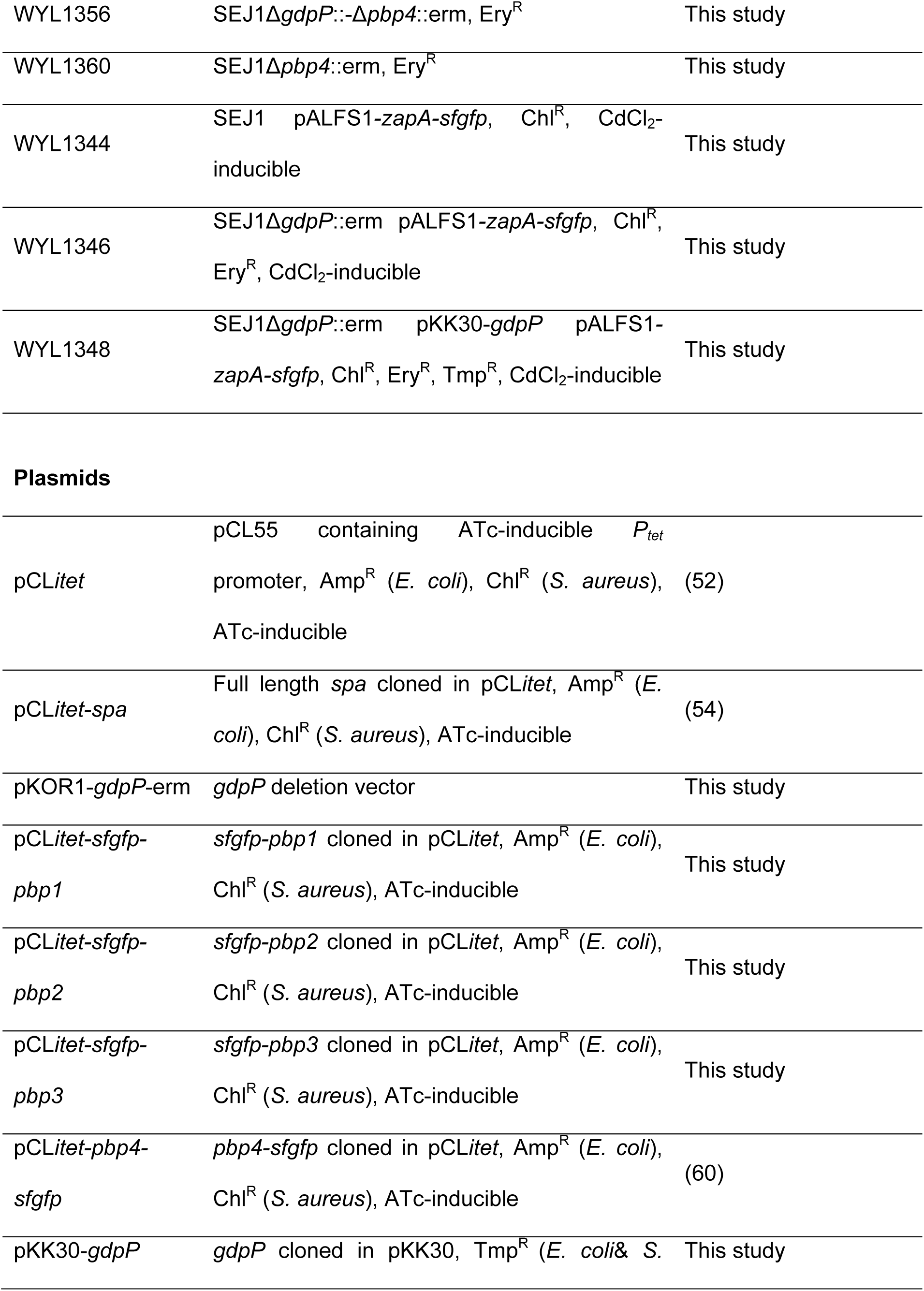
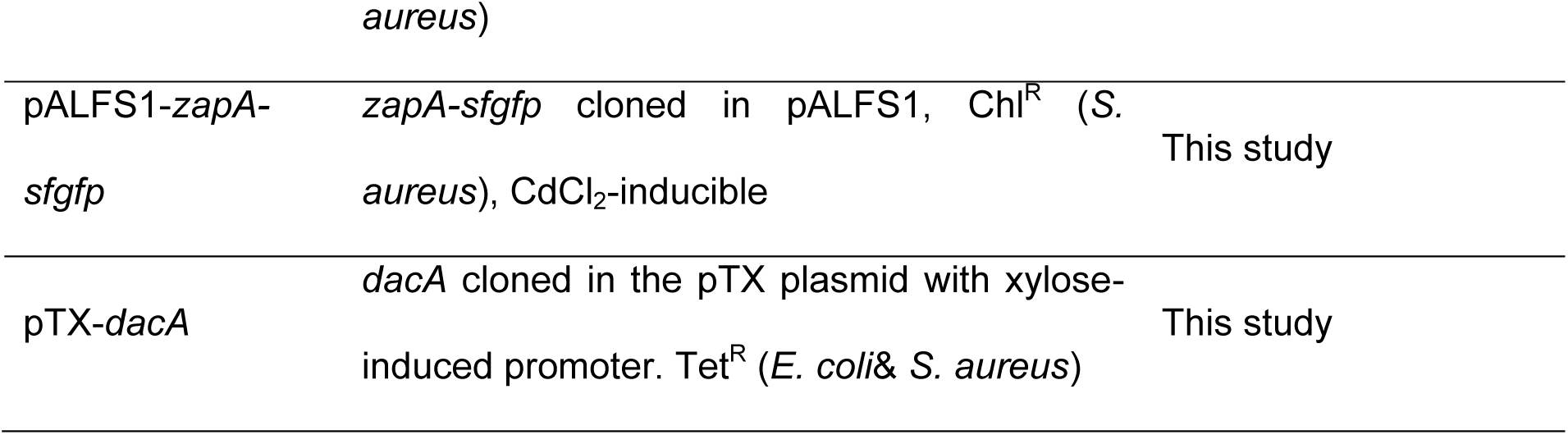
Strains and plasmids used in this study.

**Table S2.**
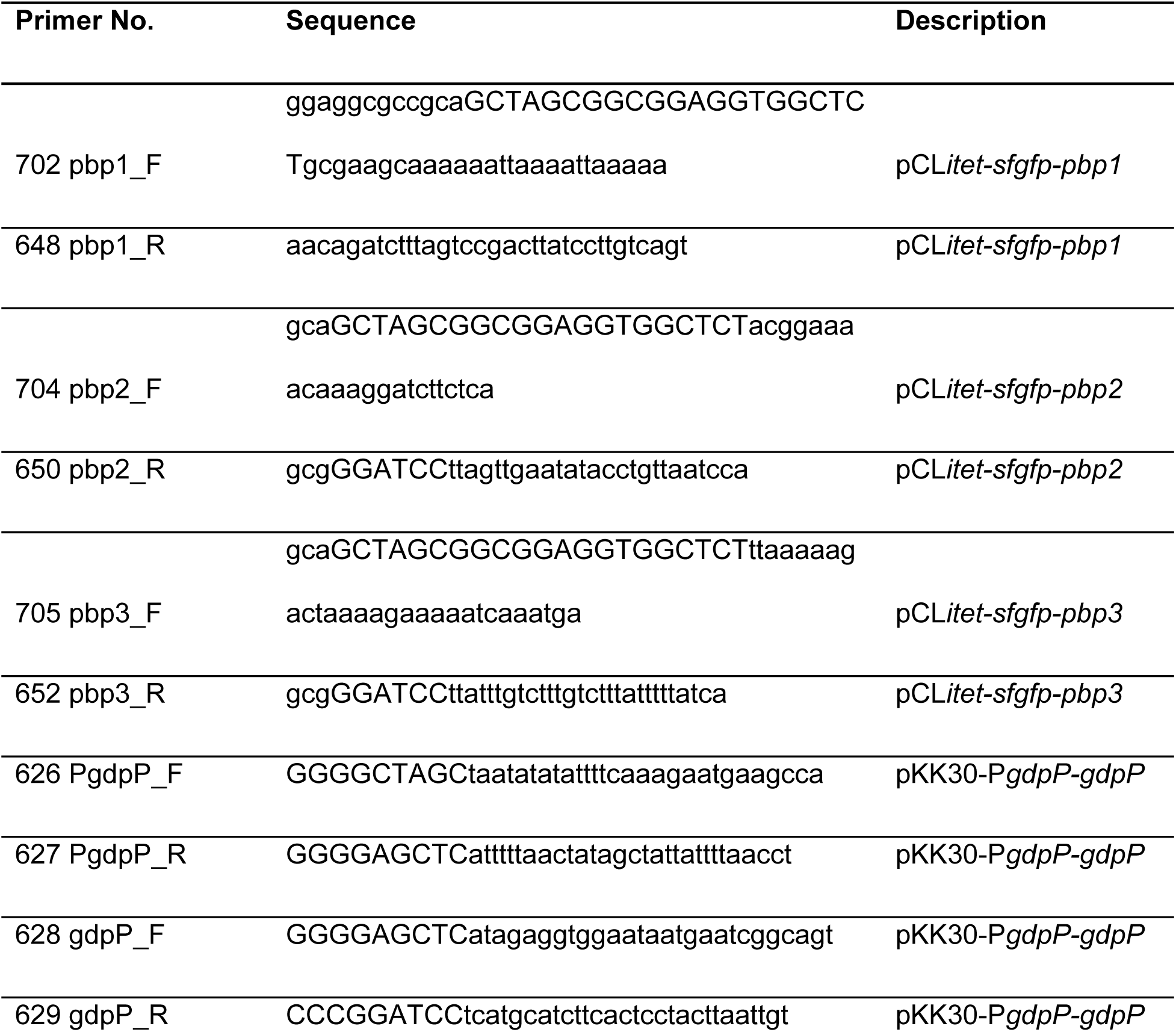
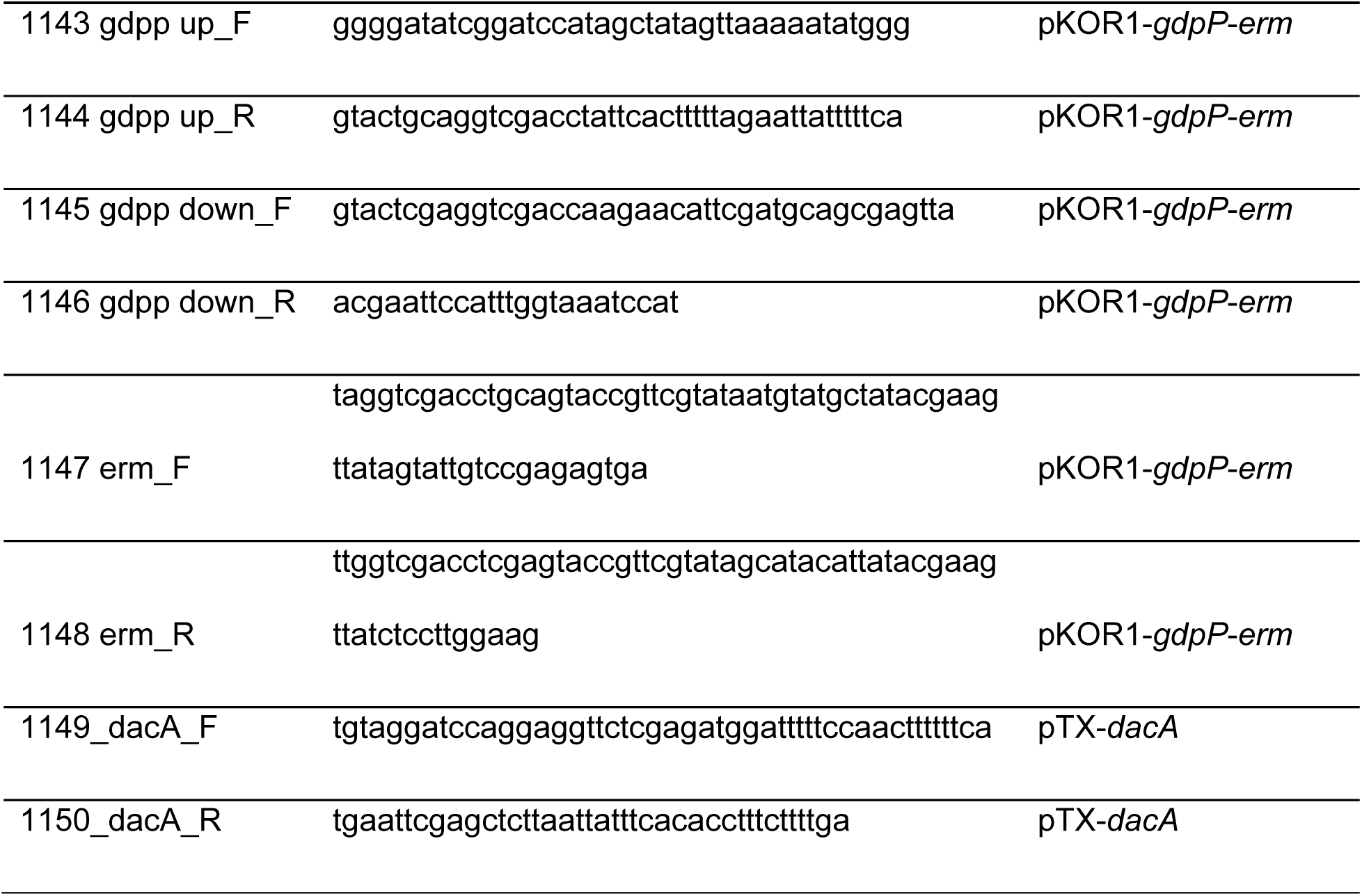
Primers used in this study.

